# Comparative genomics of *Mycoplasma feriruminatoris*, a fast-growing pathogen of wild *Caprinae*

**DOI:** 10.1101/2023.03.21.533155

**Authors:** Vincent Baby, Chloé Ambroset, Patrice Gaurivaud, Laurent Falquet, Christophe Boury, Erwan Guichoux, Joerg Jores, Carole Lartigue, Florence Tardy, Pascal Sirand-Pugnet

**Author notes:** Corresponding authors: Pascal Sirand-Pugnet Address: Université de Bordeaux, INRAE, UMR BFP, 71 avenue Edouard Bourlaux, 33140 Villenave d’Ornon, France Florence Tardy Address: Anses–Laboratoire de Lyon, VetAgro Sup, UMR Mycoplasmoses animales, 31 avenue Tony Garnier, 69007 Lyon, France. These authors contributed equally to this work (Florence Tardy, Pascal Sirand-Pugnet).

## Abstract

*Mycoplasma feriruminatoris* is a fast-growing *Mycoplasma* species isolated from wild *Caprinae* and first described in 2013. *M. feriruminatoris* isolates have been associated with arthritis, keratoconjunctivitis, pneumonia and septicemia, but were also recovered from apparently healthy animals. To better understand what defines this species, we performed a genomic survey on 14 strains collected from free-ranging or zoo-housed animals between 1987 and 2017. The average chromosome size of the *M. feriruminatoris* strains was 1,040 ± 0,024 kbp, with 24% G+C and 852 ± 31 CDS. The core genome and pan-genome of the *M. feriruminatoris* species contained 628 and 1,312 protein families, respectively. The *M. feriruminatoris* strains displayed a relatively closed pan-genome, with many features and putative virulence factors shared with species from the *M. mycoides* cluster, including the MIB-MIP Ig cleavage system, a repertoire of DUF285 surface proteins and a complete biosynthetic pathway for galactan. *M. feriruminatoris* genomes were found to be mostly syntenic, although repertoires of mobile genetic elements, including Mycoplasma Integrative and Conjugative Elements, insertion sequences, and a single plasmid varied. Phylogenetic- and gene content analyzes confirmed that *M. feriruminatoris* was closer to the *M. mycoides* cluster than to the ruminant species *M. yeatsii* and *M. putrefaciens*. Ancestral genome reconstruction showed that the emergence of the *M. feriruminatoris* species was associated with the gain of 17 gene families, some of which encode defense enzymes and surface proteins, and the loss of 25 others, some of which are involved in sugar transport and metabolism. This comparative study suggests that the *M. mycoides* cluster could be extended to include *M. feriruminatoris*. We also find evidence that the specific organization and structure of the DnaA boxes around the *oriC* of *M. feriruminatoris* may contribute to drive the remarkable fast growth of this minimal bacterium.

## Introduction

Within the prokaryotes, the class *Mollicutes* gathers bacteria characterized by their inability to synthesize peptidoglycan or the precursors necessary to build cell walls. They are consequently Gram-stain-negative, despite having evolved from Gram-positive bacteria, of which they constitute a distinct phylogenetic lineage. The class *Mollicutes* includes 11 genera (Brown et al., 2018), among which the *Mycoplasma* genus gathers the largest number of pathogenic or opportunistic species (n=78) and continues to grow as new species are regularly described in various hosts. Six new *Mycoplasma* species were described in the year 2022 alone (Noll et al., 2022; Spergser et al., 2022; Volokhov et al., 2022). *Mycoplasma* spp. genomes are small, with length varying from 580 to 1,350 kbp, resulting in very limited metabolic pathways and thus fastidious growth that generally requires sterols and complex media. Their generation time varies widely but can exceed several hours for certain species. Their G+C content is low, varying from 23% for *Mycoplasma capricolum* subsp*. capricolum* to 40% for *M. pneumoniae*. They also share a specific pattern of codon usage with UGA encoding tryptophan.

The *Mycoplasma* genus is polyphyletic and can be divided into 3 distinct groups, i.e. the two clades Hominis and Pneumoniae, and a third one known as the clade Spiroplasma. This last one includes the ‘*M. mycoides* cluster’ that contains the type species of the genus despite its eccentric phylogenetic position (Brown et al., 2018). The *M. mycoides* cluster evolved from insect-associated Mollicutes (*Spiroplasma, Entomoplasma* and *Mesoplasma*) to become ruminant pathogens capable of non-vectored direct transmission (Gasparich et al., 2004). This evolution resulted from a combination of gene losses and more than 100 novel genes gained through horizontal gene transfer from donors potentially belonging to the Hominis/Pneumoniae lineages (Lo et al., 2018). In particular, massive genetic exchanges have been predicted with the ruminant pathogen *M. agalactiae* from the Hominis clade (Sirand-Pugnet et al., 2007). Lo et al. (2018) considered species of the *M. mycoides* cluster as hybrids ‘carved’ into shared ecological niches facilitating horizontal gene transfer.

The *M. mycoides* cluster - in its strict definition, which excludes some relatively close species such as *M. yeatsii* or *M. putrefaciens* - is an ecologically, phenotypically and genetically cohesive group of five major pathogenic ruminant (sub)species whose taxonomy was amended in 2009 despite conflict between phylogeny and taxonomy (Cottew et al., 1987; Manso-Silván et al., 2007). The *M. mycoides* cluster includes four subspecies responsible for diseases listed by the World Organization for Animal Health (WOAH), namely *M. mycoides* subsp. *mycoides* (*Mmm*) and *M. capricolum* subsp. *capripneumoniae* (*Mccp*) which are the causative agents of contagious bovine and caprine pleuropneumonia, respectively, and *M. mycoides* subsp. *capri* (*Mmc*) and *M. capricolum* subsp. *capricolum (Mcap)* which are etiological agents of contagious agalactia. It also includes a fifth taxon pathogenic to cattle, *M. leachii*, which is a chimera between *mycoides* and *capricolum* species that is seldom isolated (Manso-Silván et al., 2009; Tardy et al., 2009; Fischer et al., 2012). The closely related species *M. feriruminatoris* was described ten years ago (Jores et al., 2013) and later proposed to be part of the *M. mycoides* cluster in its enlarged definition (Ambroset et al., 2017).

Over time, isolates of *M. feriruminatoris* have been collected from wild *Caprinae*, i.e. the Alpine ibex (*Capra ibex*) or Rocky Mountain goat (*Oreamnos americanus*), either in the wild or in zoos (Fischer et al., 2012, Tardy et al., 2012; Jores et al., 2013; Ambroset et al., 2017). They all share rapid growth *in vitro*, with a generation time of 27–29 min at 37°C (Fischer et al., 2013; Jores et al., 2013). Their genetic diversity was originally thought to be low, when only five isolates, mainly from a German zoo, had been investigated (Fischer et al., 2012), but was later shown to be higher once French isolates from ibex were investigated on top of the German isolates (Ambroset et al., 2017).

Despite a few dedicated papers and the availability of the genome of the type strain (G5847^T^), there are still gaps in knowledge of the *M. feriruminatoris* species. First, no specific virulence factors have been highlighted in the first genome announcement (Fischer et al., 2013), even though *M. feriruminatoris* strains are genetically equipped to produce H2O2 (Jores et al., 2013), which is a potential virulence factor of mycoplasmas, that is controversially discussed (Vilei and Frey, 2001, Szczepanek et al., 2014, Schumacher et al., 2019, Jores et al. 2020). Furthermore, *M. feriruminatoris* has been shown to produce one or two—depending on the isolate—types of capsular polysaccharides (galactan and/or β-1→6-glucan) (Ambroset et al., 2017), which is a true virulence factor for *Mycoplasma mycoides* (Gaurivaud et al., 2014, 2016; Jores et al., 2019). Second, questions remain about the level of divergence of *M. feriruminatoris* from members of the Mycoides cluster in terms of gene content and genome organization. Third, the fast-growing capacity of *M. feriruminatoris* makes it an attractive species for the rational design of vaccine chassis (Talenton et al., 2022), but the genetic bases for its fast growth are still unknown.

Here we used comparative genomics data to characterize the species *M. feriruminatoris*. The genomes of 14 *M. feriruminatoris* strains isolated over a 30-year period from 1987 until 2017, either from captive wild ruminants in zoos (n=5) or from free-roaming wild ruminants (n=9, mainly the French Alps) were compared to each other and to genomes from closely-related species in terms of synteny, gene content, phylogeny, and specific features associated with virulence or host adaptation.

## Material and Methods

### Strains, culture conditions, and molecular biology methods

*M. feriruminatoris* strains were sourced from previous studies (Fischer et al., 2012; Tardy et al., 2012; Jores et al., 2013), from the Vigimyc network (strain F11561; Poumarat et al., 2014) and isolated from the diagnostic unit of the Institute of Veterinary Bacteriology at the University of Bern (isolate 14/OD_0492). Species assignment was verified using the species-specific PCR reported previously (Ambroset et al., 2017).

All strains were cultured in PPLO medium supplemented as previously described (Poumarat et al., 1992) at 37°C with 5% CO2. Genomic DNA extraction was performed on cultures in mid-exponential phase using either 2 mL cultures for the phenol-chloroform method (Chen and Kuo, 1993) or 10 mL cultures for the NucleoBond AXG column commercial kit (Machery-Nagel).

### Genome sequencing, assembly and annotation

The full genome sequences of strains G5813/1+2, G1650, G1705, 8756-13 and 14/OD_0492 were obtained using PacBio sequencing technology, and deposited in GenBank under the accession numbers LR738858.1, LR739234.1, LR739233.1, LR739235.1, and LR739237.1, respectively. Whole-genome sequencing of *M. feriruminatoris* strains F11561, L13461, L14815, L14822, L15181, L15220, L15407 and L15568 was performed using a combination of Oxford Nanopore (ONT) and Illumina (paired-end 250 bp library) technologies (Table S1). The ONT reads were base-called using Guppy and demultiplexed using qcat (v.1.0.3, available at https://github.com/nanoporetech/qcat). The Illumina reads were trimmed using Trimmomatic (v.0.36, available at https://github.com/usadellab/Trimmomatic) (Bolger et al., 2014), and the Illumina adapters, i.e. the first 5 bp and both ends, were removed using a 5-bp sliding window with a phred score of under 20. Quality of the Illumina reads was assessed before and after trimming using fastqc (v.0.11.5, available at https://www.bioinformatics.babraham.ac.uk/projects/fastqc). The ONT reads were filtered using filtlong (v.0.2.0, available at https://github.com/rrwick/Filtlong), and reads with a length <250 bp or sharing <87% identity with the trimmed Illumina reads were excluded. The long reads were assembled using canu (v1.8, available at https://github.com/marbl/canu) (Koren et al., 2017) with an expected genome size of 1 Mbp.

The initial assembly was polished by iterative alignment of the trimmed Illumina reads using bwa-mem (v.0.7.15, available at https://github.com/lh3/bwa) (Li and Durbin, 2009) followed by correction using pilon (v.1.22, available at https://github.com/broadinstitute/pilon) (Walker et al., 2014). The final manual polishing was done by iteratively performing a variant calling pipeline and correcting the variant positions in the assemblies between each iteration. The variant calling pipeline used bwa-mem to map the Illumina reads. GATK (v.3.7, available at https://github.com/broadinstitute/gatk/) IndelRealigner (McKenna et al., 2010) was then used to perform local realignment around indels, read mate coordinates were added using the SAMtools (v.1.5, available at https://github.com/samtools/samtools) (Li et al., 2009) fixmate command, duplicate reads were marked with the Picard toolkit (v.2.18.9, available at https://broadinstitute.github.io/picard), and finally the variant positions were called using GATK HaplotypeCaller with ploidy set to 1. The corrected genomes were then manually circularized and annotated using prokka (v.1.12, available at https://github.com/tseemann/prokka) (Seemann, 2014). The completed genomes were submitted to GenBank (Table 1).

**Table 1.** *M. feriruminatoris* strains used in this study. * Plasmid accession number: CP104010

| Strain name | Host | Year isolated | Location of isolation | Main clinical sign | Genome size (bp) | Genes | CDS | IS3 | MICE | Plasmid | Accession no. |
| --- | --- | --- | --- | --- | --- | --- | --- | --- | --- | --- | --- |
| 8756-13 | Rocky Mountain goat | 1987 | USA | - | 1 060 955 | 919 | 882 | 13 | - | - | LR739235 |
| G5847<sup>T</sup> | Ibex | 1993 | Berlin Zoo, Germany | arthritis | 1 075 604 | 916 | 878 | 12 | - | - | CP091032.1 |
| G5813/1+2 | Ibex | 1993 | Berlin Zoo, Germany | arthritis | 1 037 206 | 919 | 882 | 14 | - | - | LR738858 |
| G1650 | Ibex | 1993 | Berlin Zoo, Germany | arthritis | 1 038 690 | 913 | 876 | 14 | - | - | LR739234 |
| G1705 | Ibex | 1993 | Berlin Zoo, Germany | arthritis | 1 038 707 | 915 | 878 | 14 | - | - | LR739233 |
| L13461 | Ibex | 2003 | Savoie region, French Alps | septicemia | 1 017 763 | 900 | 863 | 12 | - | - | CP113496 |
| L14822 | Ibex | 2007 | Savoie region, French Alps | pneumonia | 1 084 342 | 909 | 872 | 3 | 2 | - | CP104008 |
| L14815 | Ibex | 2007 | Savoie region, French Alps | keratoconjunctivitis | 1 021 626 | 863 | 826 | 2 | - | - | CP113497 |
| L15181 | Ibex | 2008 | Savoie region, French Alps | keratoconjunctivitis | 1 042 812 | 886 | 849 | 3 | 1 | - | CP113498 |
| L15220 | Ibex | 2009 | Savoie region, French Alps | pneumonia | 1 016 817 | 858 | 821 | 2 | 1 | 1* | CP104009 |
| L15407 | Ibex | 2010 | Savoie region, French Alps | pneumonia, keratoconjunctivitis | 1 025 568 | 966 | 929 | 2 | - | - | CP113499 |
| L15568 | Ibex | 2011 | Savoie region, French Alps | none (animal follow-up) | 1 015 346 | 950 | 913 | 1 | - | - | CP113500 |
| 14/OD_0492 | Ibex | 2014 | Nature and Animal Park Goldau, Switzerland | liver, fibrinous polyarthritis | 1 070 562 | 941 | 904 | 0 | 1 | - | LR739237 |
| F11561 | Ibex | 2017 | Haute-Savoie region, French Alps | Unknown (animal found dead in the wild) | 1 011 470 | 854 | 817 | 1 | - | - | CP113495 |

### Comparative genomic analysis

Homologous protein sequence clustering was performed using get_homologues (v.05032019, available at https://github.com/eead-csic-compbio/get_homologues) (Contreras-Moreira and Vinuesa, 2013) with the COGtriangle (Kristensen et al., 2010) clustering algorithm. Multiple sets of genomes were analyzed. One set included the *M. feriruminatoris* genomes only and was used to identify the core genome and pan-genome of this species. A second set included the *M. feriruminatoris* genomes as well as the *Mmc* GM12 genome (RefSeq accession no. NZ_CP001668) and was used to build the *M. feriruminatoris* phylogenetic tree. A third set included the *M. feriruminatoris* genomes, multiple complete genomes from members of the *M. mycoides* cluster, i.e. *Mmc* GM12 and 95010 (RefSeq accession nos. NZ_CP001668 and NC_015431), *Mmm* Glasdysdale, PG1, Ben1, Ben50, Ben468, izsam_mm5713 and T1/44 (RefSeq accession nos. NC_021025, NC_005364, NZ_CP011260, NZ_CP011261, NZ_CP011264, NZ_CP010267 and NZ_CP014346), *Mcap* ATCC 27343 (RefSeq accession nos. NC_007633) *Mccp* 9231-Abomsa, 87001, M1601, ILRI181 and F38 (RefSeq accession nos. NZ_LM995445, NZ_CP006959, NZ_CP017125, NZ_LN515399 and NZ_LN515398) and *M. leachii* PG50 (RefSeq accession no. NC_014751). This set also included other *Mycoplasma* strains, i.e. *M. putrefaciens* KS1, Mput9231 and NCTC10155 (RefSeQ accession no. NC_015946, NC_021083 and NZ_LS991954), *M. yeatsii* GM274B (RefSeq accession no. NZ_CP007520). Finally, the genome of *Mesoplasma florum* L1 (RefSeq accession no. NC_006055) was also used as an outgroup for construction of the phylogenetic tree.

In order to compare core and pan-genomes of *M. feriruminatoris* and related species, all the proteins of a combined set including the *M. mycoides* cluster members, *M. yeatsii*, *M. putrefaciens, Mesoplasma (Me) florum and M. feriruminatoris* were clustered based on sequence similarity. Three sets of genomes were compared, each with their own core and pan-genome reconstructed based on these clusters. One set was formed with the 14 *M. feriruminatoris* strains, the second set was formed with the 4 *M. yeatsii* and *M. putrefaciens* strains, and the third set was formed with the 16 *M. mycoides*-cluster strains available in databases. *Me. florum* was excluded from the sets, but its proteins were used as an outgroup during the clustering process.

Synteny of the *M. feriruminatoris* genome was analyzed by whole-genome alignment using Mauve (v.20150226, available at https://darlinglab.org/mauve/mauve.html) (Darling et al., 2004, 2010). Synteny of the single-copy core genes in all the *Mycoplasma* genomes was visualized using the GMV genome browser (v.1e-93, available at http://murasaki.dna.bio.keio.ac.jp/wiki/index.php?GMV) (Popendorf et al., 2010). Proteins located in the putative *M. feriruminatoris* Mycoplasma Integrative Conjugative Elements (MICE) were compared to the proteins of ICEA5632-I from *M. agalactiae* 5632 and ICEM from *Mmc* GM12 (Citti et al., 2018) using BLASTp (v. 2.9.0, available at https://ftp.ncbi.nlm.nih.gov/blast/executables/blast+/LATEST/). Insertion sequence (IS) detection was performed using ISfinder (available at https://isfinder.biotoul.fr/) (Siguier et al., 2006).

### DUF285 protein analysis

All predicted protein sequences in the *M. feriruminatoris* genomes were screened for DUF285 domains using CD-search (Marchler-Bauer and Bryant, 2004). Motif detection was then performed using MEME (v. 5.0.5, available at https://meme-suite.org/meme/) (Bailey et al., 2009), initially with default parameters but then adjusting the motif length to 25 and 16 amino acids in two separate runs to refine the motifs. The motifs found in all the predicted proteins were then detected using MAST (v. 5.0.5, available at https://meme-suite.org/meme/) (Bailey et al., 2009) in its default parameters. Protein signal peptides were predicted using SignalP (v. 6.0, available at https://services.healthtech.dtu.dk/services/SignalP-6.0/) (Teufel et al., 2022), and transmembrane domains were predicted using DeepTMHMM (available at https://dtu.biolib.com/DeepTMHMM) (Hallgren et al., 2022).

### *In silico* analysis of polysaccharide pathways

tBlastX analysis was used to retrieve putative enzymes involved in polysaccharide synthesis and described in *M. feriruminatoris* strain G5847^T^ (Ambroset et al., 2017) and in other mycoplasma species (Gaurivaud et al., 2016; Schieck et al., 2016). A prediction of transmembrane regions was carried out by TMHMM2 (available at https://services.healthtech.dtu.dk/services/TMHMM-2.0/)(Krogh et al., 2001) and the synthase-specific cytoplasmic domain with the DXD and R/QXXRW-like motifs were identified by alignment with the galactan synthase MSC_0108 from *Mmm* PG1^T^ or GsmA from *M*. *agalactiae* 14628.

### Phylogenetic tree construction

The phylogenetic trees were constructed using respectively 555 single-copy core genes for the intra-species tree (Figure 1B) and 294 single-copy core genes for the tree including other strains for the *M. mycoides* cluster (Figure 5). For each set, the protein sequences were aligned using Clustal Omega (v.1.2.1, available at http://www.clustal.org/omega/) (Sievers et al., 2011) and the alignments were then concatenated. Unaligned and low-confidence regions were removed from the alignment using Gblocks (v.0.91b, available at molevol.cmima.csic.es/castresana/Gblocks.html) (Talavera and Castresana, 2007), thus producing sequence matrices of 186,920 and 92,785 amino acid sites for the *M. feriruminatoris* tree and the *M. mycoides* tree, respectively. The evolution model for tree construction was determined using ProtTest (v.3.4.2, available at https://github.com/ddarriba/prottest3) (Darriba et al., 2011), and in both cases the CpREV model (Adachi et al., 2000) was identified as the best model. The trees were then created with RaxML (v.8.2.12, available at https://github.com/stamatak/standard-RAxML) (Stamatakis, 2014) using the GAMMA model of rate of heterogeneity, and 450 and 150 bootstrap replicates were made for the *M. feriruminatoris* tree and the *M. mycoides* tree, respectively, using the autoFC bootstopping criterion to determine the number of replicates.

**Figure 1.**
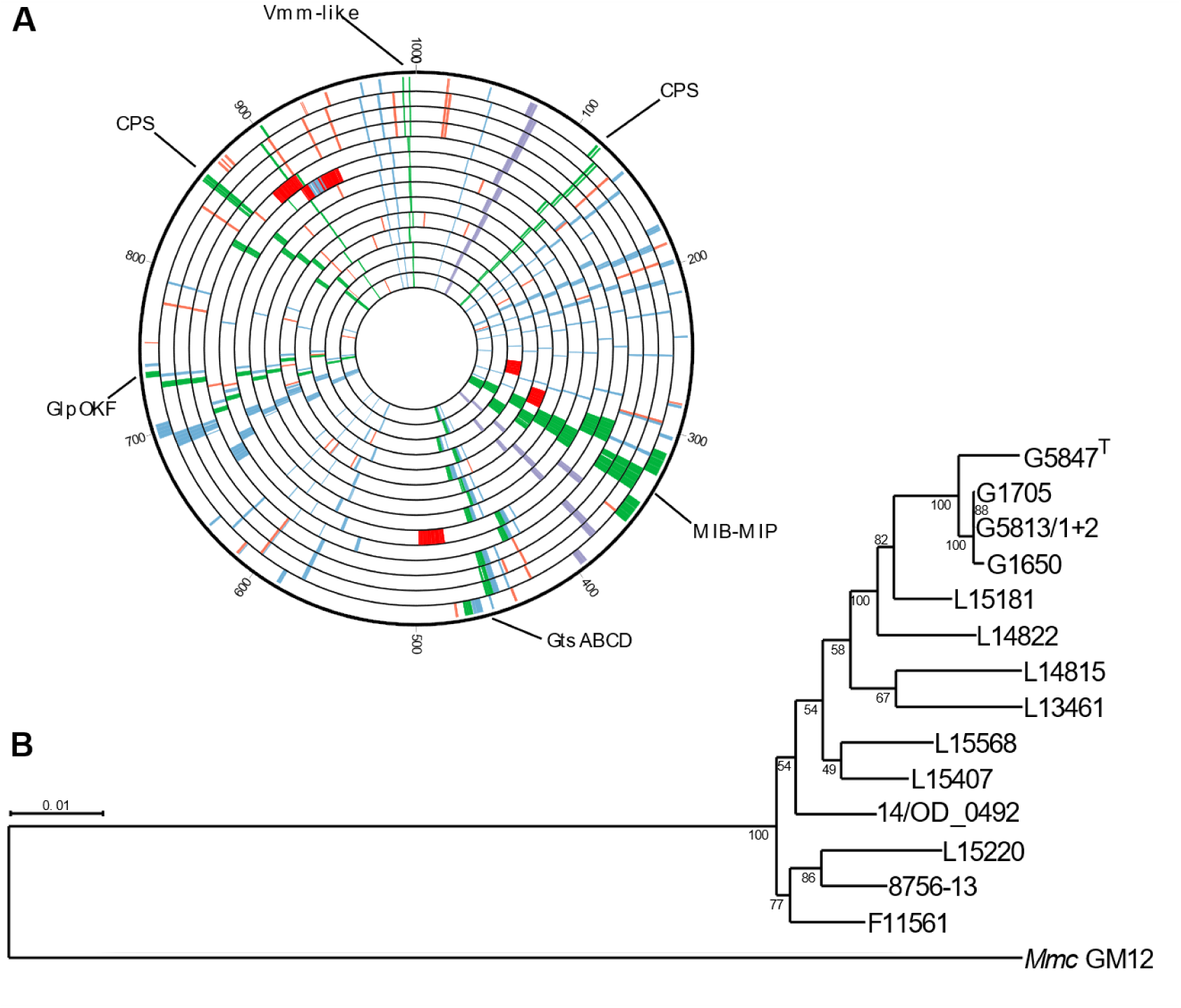
Genomic structure and phylogeny of the *M. feriruminatoris* genomes. **(A)** circular representation of all the *M. feriruminatoris* genomes, with size normalized to 1 Mbp. From outer to inner circles, the genomes are G5847^T^, G1650, G1705, G5813/1+2, L15181, L14822, L14815, L15407, L15568, 14/OD0492, L15220, 8756-C13, F11561 and L13461. Genes are colored based on function, with virulence genes in green, insertion sequences (IS) in orange, rRNAs in purple, DUF285 proteins in blue, and MICE genes in red. **(B)** Phylogenetic tree of the *M. feriruminatoris* strains. The tree was constructed using the maximum-likelihood method and inferred from the concatenated alignments of 555 single-copy core protein sequences. The alignment matrix contained 186,920 amino acid sites. *Mmc* strain GM12 was used as the outgroup, and 450 bootstrap replicates were run. Bootstrap values are shown as node labels.

### Ancestral genome reconstruction

The evolution of the gene-family contents over the course of the evolution of *Mycoplasma* species included in this work was studied using the COUNT software (available at http://www.iro.umontreal.ca/∼csuros/gene_content/count.html) (Csurös, 2010). We used the phylogenetic tree described above with 14 *M. feriruminatoris* strains, 16 M. *mycoides* cluster-related strains, 3 *M. putrefaciens* strains, 1 *M. yeatsii* strain and the outgroup *Me. florum*, and we used the presence/absence matrix produced in the comparative analysis to monitor the occurrence of 2615 gene families. We used a birth-and-death model to calculate the posterior probabilities, and we used a gain–loss model with a Poisson distribution at the root and set the edge length, loss and gain rates at 4 gamma categories to maximize the likelihood of the optimized model.

## Results and Discussion

### A homogeneous species with a closed pan-genome

In order to get an overview of the *M. feriruminatoris* species and further investigate its evolutionary relationship with other species from the *M. mycoides* cluster, we sequenced the genome of 13 strains isolated from different regions and years and from animals showing different clinical signs (Table 1). We used a combination of Illumina short reads and PacBio or ONT long reads to produce complete circular chromosome sequences for all the strains. Strain G5847^T^ was included in the comparative genomic analysis. The average chromosome size of the 14 strains was 1,040 ± 24 kbp with the smallest and largest genome at 1,011,470 bp and 1,084,342 bp for strains F11561 and L14822, respectively (Table 1). As expected for members of the class *Mollicutes*, the G+C content of the genomes was low, at an average of 24.24% ± 0.04%. The genomes were annotated using prokka (Seemann, 2014) that predicted between 853 and 941 (average 890) genes per genome, including 816 to 904 (average 852) protein-encoding genes (Table 1). Each genome had two rRNA loci encoding the 5S, 16S and 23S rRNAs separated by approximately 318 kbp and located on the same half of the genome relative to the chromosomal origin of replication and the terminus (Figure 1A). A total of 30 tRNAs genes and a single tmRNA were predicted in every genome.

A total of 11,935 proteins were predicted from the 14 *M. feriruminatoris* genomes. A phylogenetic tree was built using the sequence of 555 single-copy core proteins found in all *M. feriruminatoris* strains and in *Mmc* strain GM12 which was used as the outgroup (Figure 1B). In the resulting tree, *M. feriruminatoris* strains were grouped into a homogeneous single branch at a short distance from the *Mmc* root. Four strains, i.e. G5847^T^, G1650, G1705 and G5812/1+2, that had been isolated in a narrow time-window (1993–1994) from ibexes that were hosted in Berlin zoo and showed similar clinical signs, were very closely related, as expected. However, besides these zoo strains, we found no further correlation between phylogenetic branches and location or time of isolation. For example, the most-recently isolated strain F11561 (2017) and the least-recently isolated strain 8756-13 (1987), which were also isolated on two separate continents (Europe and North America, respectively), were found in the same branch. The core and pan-genomes of the *M. feriruminatoris* species were determined based on the 11,390 proteins grouped in 1,312 clusters of homologs retrieved from the 14 *M. feriruminatoris* genomes (Figure 2).

**Figure 2.**
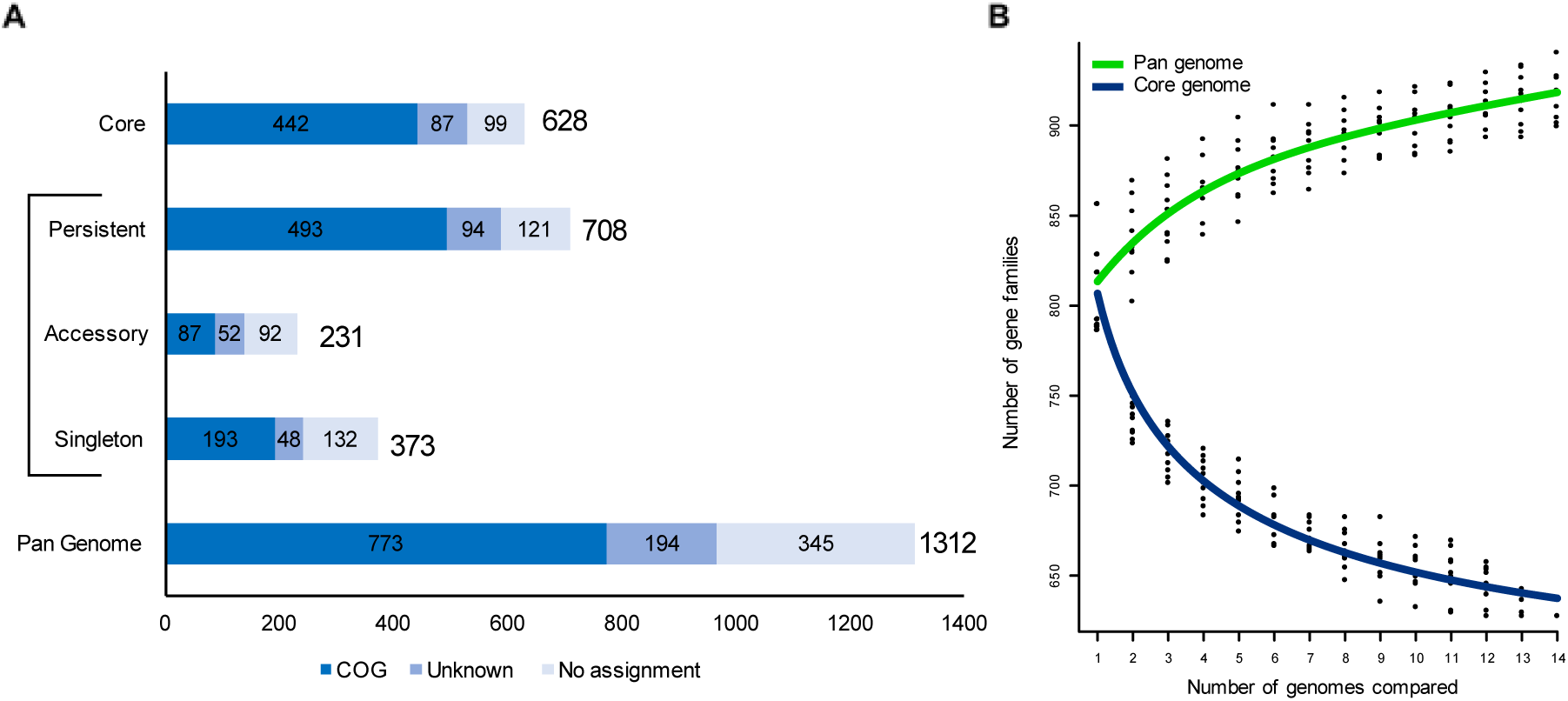
Core and pan-genomes of the *M. feriruminatoris* species. **(A).** The pan-genome is composed of 1,312 gene families encoding proteins, of which 373 are singletons (present in a single genome), 708 are persistent (present in at least 13/14 genomes), 628 are core (present in all genomes), and the remainder (n= 301) are accessory (present in 2 to 12 genomes). The sum of persistent, accessory and singleton genes constitute the total pan-genome genes. Gene families with predicted eggNOG (NOG) function categories are indicated as follows: dark blue, known; medium blue, unknown; light blue, no assignment. (**B).** Gene number estimation curves for the *M. feriruminatoris* core genome (blue, bottom curve) and pan-genomes (green, top curve) were generated using the methods described in Willenbrock et al. (2007) and Tettelin et al. (2005), respectively.

A core set of 628 clusters of protein-encoding genes was present in every genome, representing an average of 47.9% of all clusters within a given strain. Considering possible sequencing errors, this value was extended to 708 clusters of persistent gene predicted from at least 13 out of the 14 genomes.

A total of 373 singleton protein clusters representing 28.4% of all clusters were present in only one strain. The *M. feriruminatoris* strains had an average of 27 ± 17 singleton clusters representing 3.2% of their total number of different clusters, but most of them contained short protein sequences measuring less than 200 amino acids, which suggests that many could be annotation artifacts (i.e. pseudogene remnants). These potential artifacts in strain-specific proteins together with the slow rise of the pan-genome estimation curve (Figure 2B) suggest that the *M. feriruminatoris* species possesses a closed pan-genome with most gene families shared between multiple strains, which is consistent with the recently proposed genomic definition of a bacterial species (Moldovan and Gelfand, 2018).

A detailed analysis of the distribution of gene families from the core, persistent, accessory and singleton genomes into different functional categories was then undertaken (Figure S1). Among the 1312 gene families of the *M. feriruminatoris* pan-genome, only 773 (58.9%) were assigned to non-supervised orthologous groups (eggNOG) with one or several functional categories, whereas 194 (14.8%) were assigned to an eggNOG with unknown functional category, and 345 (26.3%) were not assigned. In the *M. feriruminatoris* core and persistent genomes, there were 444 (33.8%) and 506 (38.6%) clusters assigned to an eggNOG with a known function, respectively, most of which were related to genetic information storage and processing (14.2% of core and 16.2% of persistent genomes) and metabolism (14.9% of core and 16.2% of persistent genomes). Only 89 (6.9%) clusters from the accessory genome were assigned to an eggNOG with a known function, of which 41 (3.1%) were associated with genetic information storage and processing functions. Within the singleton genome, 132 (10.1%) clusters were not assigned to an eggNOG, and 48 (3.6%) were assigned to an eggNOG with an unknown function. Of the 193 clusters (51.7% of singletons) with known eggNOG categories, 106 (28.7% of singletons) were related to genetic information storage and processing. Strain-specific clusters included a substantial proportion of proteins dedicated to defense mechanisms (V category, 22/373 against 13/628 in the core genome) (Figure S1). Although the eggNOG domain of ‘genetic information storage and processing’ categories appeared to be highly represented in core, persistent, accessory and singleton genomes, further analysis revealed significant differences. Although the persistent genome encompassed 82.3% (126/153) and 66.6% (32/48) of gene families from categories J (translation, ribosomal structure and biogenesis) and K (transcription), it only encompassed 35.5% (54/152) of gene families for category L (replication, recombination and repair), whereas the singleton genome encompassed 41.5% (63/152) of category-L gene families, which suggests fast turnover of some genes. Further analysis indicated that many of these highly volatile genes could be associated with mobile genetic elements (MGE) such as IS or MICEs. The 154 gene families involved in cellular processes and signaling represented 17.8% of the gene families with known eggNOG function categories, the most represented category being V (defense mechanism) with 16 and 22 families present in accessory and singleton genomes, respectively, which also suggests a fast evolution of the corresponding repertoire of genes among *M. feriruminatoris* strains. Taken together, our findings from detailed analysis of the *M. feriruminatoris* pan-genome point to a global conservation of genes families involved in information processing and central metabolism and more strain-specific repertoires associated with MGEs and defense mechanisms.

### A highly syntenic structure, locally influenced by the mobilome

Synteny blocks from the 14 *M. feriruminatoris* genomes shared mostly the same order (Figure 3A). There was no observable major reorganization except one noticeable ∼35 kbp duplication event in the G5847^T^ genome (Talenton et al., 2022), resulting in six copies of the immunoglobulin cleavage proteins MIB and MIP (Arfi et al., 2016) while the other strains had either three or four copies.

**Figure 3.**
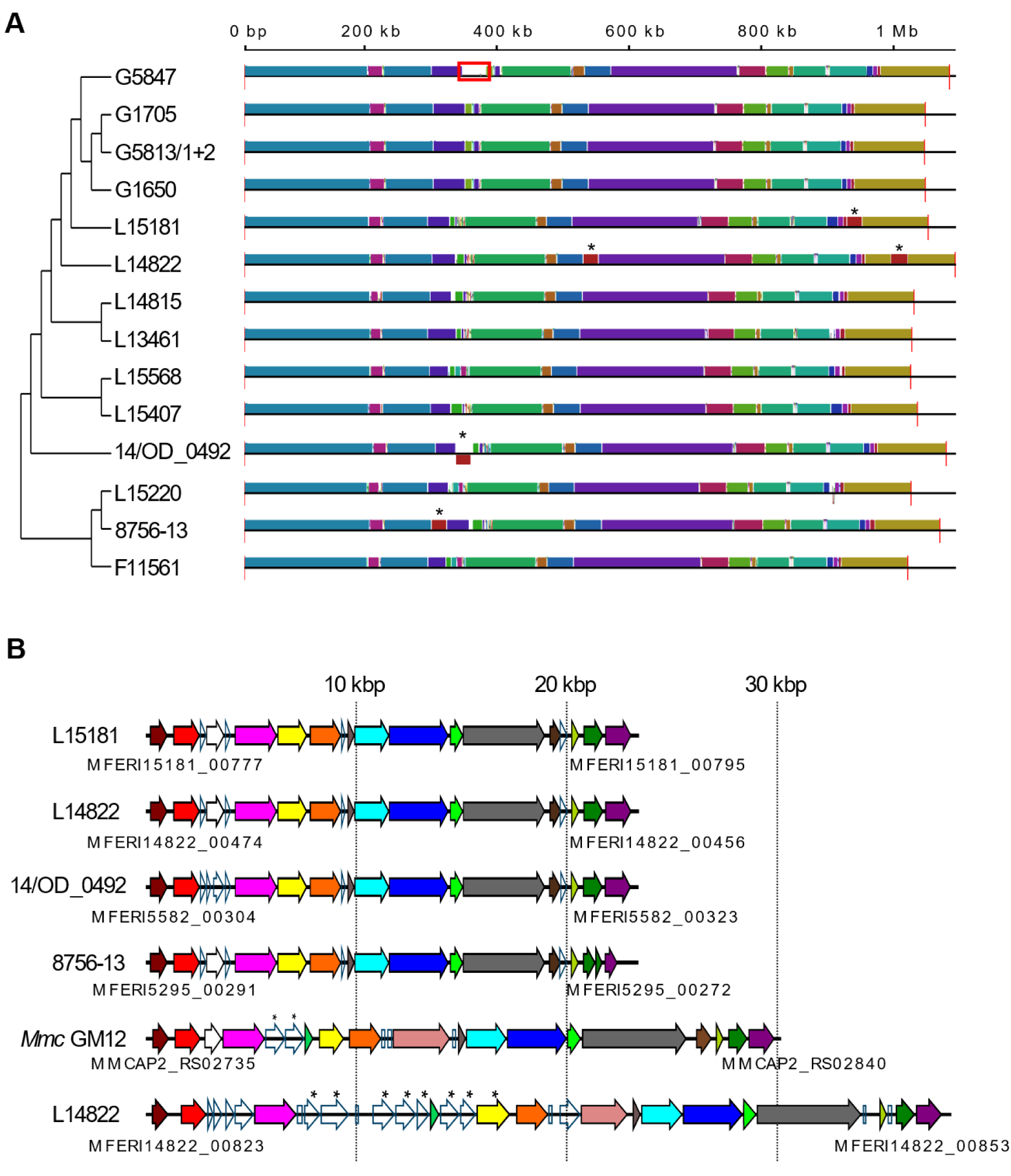
Synteny between the *M. feriruminatoris* genomes. **(A)** Conserved synteny blocks shared by the different strains. The tree topology represented on the left side corresponds to the tree in Figure 1B. MICE locations are identified with an asterisk, and the 35 kbp duplication in strain G5847T is framed in red. **(B)** *M. feriruminatoris* MICE structure. Homologous genes between the *M. feriruminatoris* and *Mmc* GM12 MICEs are represented with the same color. The MICE strain of origin is indicated on the left of each MICE, and the locus tags of the first and last genes of the MICEs are specified under their respective track.

A synteny block of ∼23 kbp found in strains L15181, L14822, 14/OD_0492 and L15220 was shown to be located in different regions and orientations depending on the strain (Figure 3A, red block tagged with an asterisk). Upon closer inspection, these loci were shown to be akin to the MICE identified in *Mmc* GM12 (Figure 3B). Almost all the *Mmc* GM12 MICE genes had a homolog in the *M. feriruminatoris* MICEs and were also in the same order. A second putative MICE was identified in strain L14822, although with a slightly different structure and encoding multiple proteins within the DUF285 domain. With the exception of strain 14/OD-0492 that had none, all *M. feriruminatoris* strains presented variable numbers of IS highly related (>87% transposase similarity) to IS3 found in *M. mycoides*-cluster mycoplasmas (Table 1). Most strains had up to three complete or degraded copies, while some, notably those from zoo isolates, had 12 to 14 copies. The surge of ISs observed in these strains did not modify the genome organization, which contrasts with large inversions evidenced in the genomes of some *M. mycoides*-cluster strains such as *Mmc* 95010 (Thiaucourt et al., 2011), PG3 and 152/93 (Hill et al., 2021) and *Mmm* T1/44 (Gourgues et al., 2016). We found no traces of prophages or CRISPR loci in *M. feriruminatoris* genomes. Only one out of the 14 strains (L15220) was shown to harbor a 3,301 bp plasmid. The pL15220 plasmid was longer than most of the plasmids described so far within the *M. mycoides* cluster but close to the size of pMyBK1 plasmid from *M. yeatsii* (3,432 bp) (Breton et al., 2012), and had a G+C content of 26.5%. The strongest pairwise nucleotide identity with mycoplasma plasmids was obtained with pMmc-95010 (1,850 pb and 48.6% of identity) (Thiaucourt et al., 2011), despite a noticeable difference in size of circa 1,400 bp.

The pL15220 plasmid encodes two CDSs, including a hypothetical protein and a predicted CopG-family transcriptional regulator sharing 82% nucleotide identity with its homolog in pMmc-95010 and a second CDS (649 aa) with no homolog in databases (Figure S2). Noticeably, three regions of 94 to 293 nt shared 76%–100% nucleotide identity with three different regions of the L14822-specific MICE, a feature already described in work comparing the plasmid and MICE of the *Mmc* 95010 strain (Thiaucourt et al., 2011) and that suggests genetic exchanges among these MGEs. The pL15220 plasmid does not encode a Rep2 protein with a pfam01719 domain shared by most replicases (Breton et al., 2012). This finding is consistent with the absence of a double-strand origin (dso) where the Rep proteins normally cleave (at a conserved site TACTAC(C)G/A) the positive DNA strand to start the rolling-circle replication (Moscoso et al., 1995). In contrast, an sso block (lagging-strand initiation site) similar to pMmc-95010 was identified next to the HP encoding gene. Its mosaic structure together with the presence of a CDS encoding a hypothetical protein with no homologs in databases suggests that the pL15220 plasmid may belong to a new plasmid family whose emergence was marked by recombination events with other mycoplasma plasmids and MICEs.

Because of its overall highly conserved genome synteny and its coherent positioning within the intraspecies phylogeny tree, *M. feriruminatoris* G5847^T^ might be considered a valuable representative of the species. In many cases, the very first strain chosen for genome sequencing is a lab strain that has encountered an unknown number of passages and whose origin often remains unclear. This was the case for the *Mmm* PG1^T^ genome sequenced in 2004 whereas the isolate dated back to 1931 (Westberg et al., 2004). Soon after, PG1^T^ was shown to differ greatly from other field strains with a notable 24-kb genetic locus repetition and hence was not the best representative of this important bovine pathogen (Bischof et al., 2006). In the case of *M. feriruminatoris*, the G5847^T^ isolate also has one major, near-perfect 35 kbp duplication that was not present in other genomes. This duplication was ascertained using two independent long-read sequencing strategies (Talenton et al., 2022), and we concluded it may have occurred during passages in the lab.

### A variable repertoire of enzymes involved in polysaccharides biosynthesis

*M. feriruminatoris* isolates were previously shown to produce a capsule composed of either galactan and/or β-(1→6)-glucan depending on the isolates (Ambroset et al., 2017). We used tBlastX to search for homologs of enzymes involved in polysaccharide biosynthesis, as predicted in *M. feriruminatoris* G5847^T^ and in other *Mycoplasma* species (Gaurivaud et al., 2016; Ambroset et al., 2017; Schieck et al., 2016). A complete putative biosynthetic pathway for galactan was predicted in all *M. feriruminatoris* isolates, with genes clustered in two different genomic locations (Figures 1 and 4).

**Figure 4:**
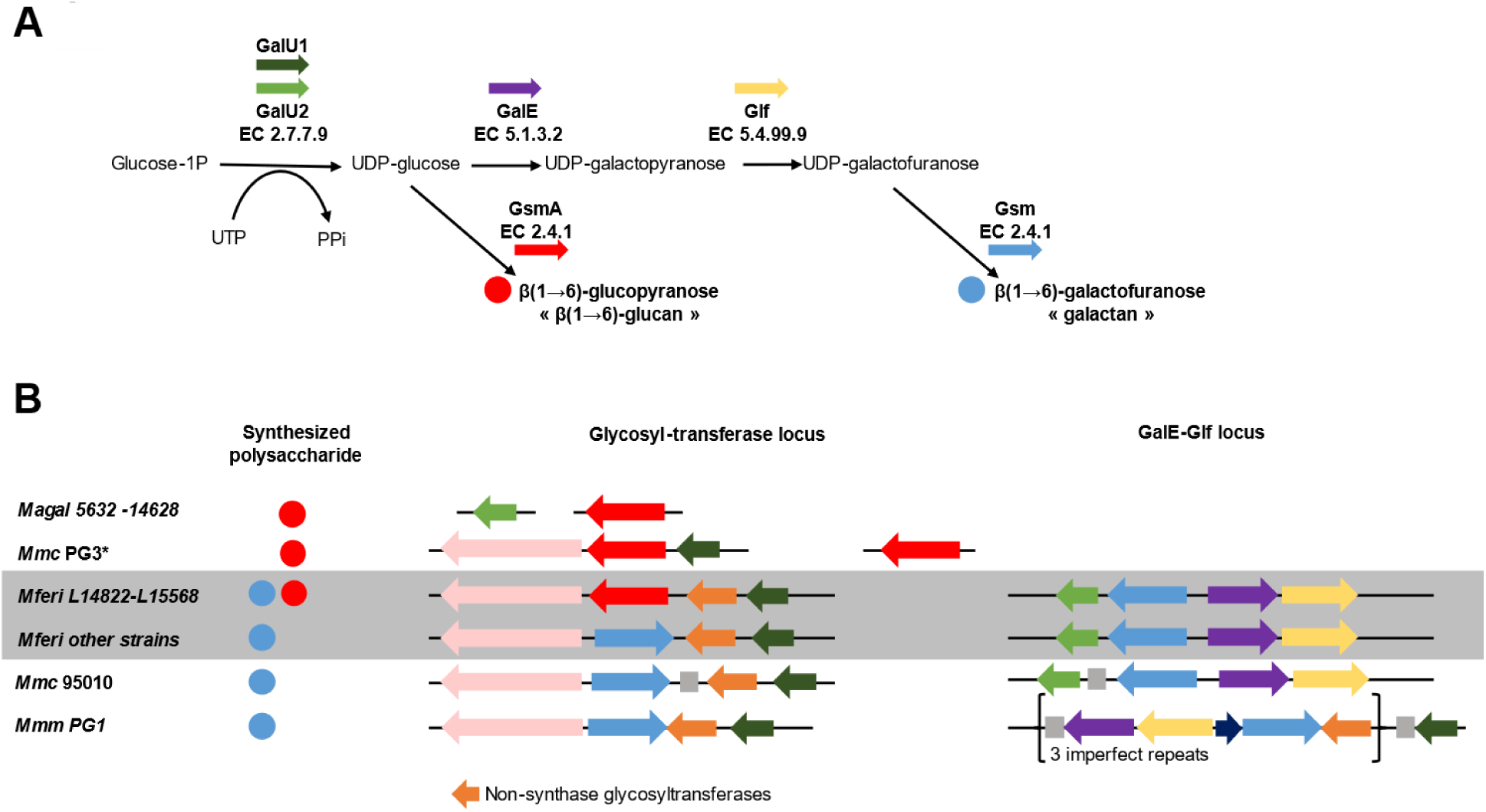
Polysaccharide biosynthesis pathways. **(A)** Schematic representation of metabolic pathways involved in β(1→6)-glucan and galactan synthesis. Glucose-1P is transformed into UDP-glucose by a glucose-1-phosphate uridylyltransferase (GalU1 or GalU2). An UDP-glucose 4-epimerase (GalE) and an UDP-galactofuranose mutase (Glf) successively transform the UDP-glucose into UDP-galactofuranose. Finally, one (or several) glycosyltransferases (GT) with synthase activity (<u>G</u>lycan <u>S</u>ynthase of <u>M</u>ollicute, Gsm or GsmA for *M. agalactiae*) builds and exports the final polysaccharide, a polymer of galactofuranose, the galactan (blue circle) or β(1→6)-glucan (red circle). **(B)** Organization of the genes encoding the corresponding enzymes amongst genomes of the *M. mycoides*-cluster strains and *M. agalactiae* (*M. agal*). Red or blue circles indicate the nature of the polysaccharide (as in panel A), colored arrows represent coding sequences (synthases are in blue or red as in panel A, other glycosyltransferases lacking the structural signature of synthases are in orange, and dark blue and pink arrows represent hypothetical proteins and peptidases, respectively). Grey squares represent transposases.

Eleven out of the thirteen newly sequenced strains carried two nearly identical *Gsm* genes encoding genuine galactan synthases that were homologs to the synthase encoded by MSC_108 in *Mmm* PG1^T^ (Gaurivaud et al., 2016) and carry the typical 4 transmembrane domains (TMDs) and a cytoplasmic domain with DAD and QRMRW motifs. One is located in the glycosyl-transferase locus, and the other is located in the GalE-Glf locus (Figure 4B). In contrast, but in agreement with previous PCR findings (Ambroset et al., 2017), isolates L14822 and L15568 have one *Gsm* copy only. However, these two isolates harbor a different synthase, at the exact same position as the first *Gsm* in other strains, with 7 TMDs and a cytoplasmic domain harboring DXD and RXXRW motifs, homologous to the GsmA synthase involved in β-(1→6)-glucan production in *M. agalactiae* 14628 and *Mmc* PG3 (Bertin et al., 2015; Gaurivaud et al., 2016).

The GsmA protein of *M. feriruminatoris* is closer to its homolog in *Mmc* PG3 than in *M. agalactiae* 14628 (81.2% vs 66.5% protein identity). The *GsmA* proteins of isolates L14822 and L15568 are 99.9% identical. This ‘surgical’ genomic ‘replacement’ of a *Gsm* gene by a *GsmA* gene in two isolates only and the mechanism beyond the replacement are an intriguing feature of the otherwise very syntenic clusters for polysaccharide biosynthesis.

### A large repertoire of lipoproteins and proteins with DUF285 domains

Using SignalP, we found 1,104 lipoprotein signal peptides in the 14 *M. feriruminatoris* genomes corresponding to an average of 79 [77–83] lipoproteins per strain. BlastP analyses confirmed the presence of the usual main immunogenic lipoprotein repertoires of the *M. mycoides* cluster, including LppA, LppB, LppC, LppQ and the variable surface protein Vmm. In each genome, LppB, LppC or LppQ genes were present in a unique copy, whereas up to four genes encoding LppA and 6 to 7 Vmm copies were detected per genome. Interestingly, isolates collected from wild ibex had 3 to 4 copies of the LppA gene, with either two pairs of adjacent genes in different regions or one pair and a singleton, whereas isolates from zoos harbored one to three copies of the gene and several (up to 4) truncated genes. The proximity of IS could explain the presence of duplicated, truncated LppA-encoding genes.

Lipoproteins with multiple DUF285 domains were also detected in all *M. feriruminatoris* genomes (Figure S3). DUF285 domains are presented as one of the top 10 relevant domains for animal host classification (Kamminga et al., 2017). Proteins with DUF285 domains, of as-yet unknown function, are also found in the members of the *M. mycoides* cluster (Sirand-Pugnet et al, 2007) and in other ruminant mycoplasmas including *M. agalactiae* and *M. bovis*. DUF285-domain proteins contain varying numbers of 25 amino acid long-tandem repeats (Röske et al, 2010). The tandem repeats of the *M. feriruminatoris* DUF285 proteins are also preceded by a 16-residue motif of 12 to 14 amino acids upstream of the first repeat (Figure S3A). This upstream motif is found for almost every DUF285 protein, the only exceptions being proteins < 200 amino acids long (Figure S3B). Furthermore, this upstream motif was only found in DUF285 proteins. A total of 380 DUF285 proteins were found in *M. feriruminatoris* genomes, i.e. an average of 27 DUF285 domain proteins per strain. The DUF285 proteins grouped into 51 clusters based on the comparative genomic analysis, and 11 of them were part of the core genome. Two of the core DUF285 protein clusters are also duplicated between two to five times in every strain. The DUF285 protein clusters present in at least 10 strains are mostly predicted to also possess a signal peptide or a lipoprotein signal peptide, but some are also predicted to have transmembrane domains at either the C or N terminus. When more than one transmembrane domain was predicted, they were located at both the C and N terminus of the proteins. Remarkably, 8 CDS encoding DUF285 proteins were all found in the L14822-specific MICE (Figure S2).

### *M. feriruminatoris* is much closer to any member of the *M. mycoides* cluster than to either *M. putrefaciens* or *M. yeatsii*

To evaluate the relatedness and specificities of *M. feriruminatoris* compared to mycoplasmas from the *M. mycoides* cluster—in its strict definition—and closely-related ruminant species (i.e. *M. yeatsii* and *M. putrefaciens*), we ran a global comparative genomics analysis including 21 genomes in addition to the 14 from *M. feriruminatoris* (Table S2). In a high-resolution phylogenetic reconstruction based on 294 single-copy core proteins, *M. feriruminatoris* appeared much closer to any of the *M. mycoides* cluster members than to *M. putrefaciens* or *M. yeatsii* (Figure 5A). This result is in accordance with previous phylogenetic studies based on 16S rRNA gene sequences (Jores et al., 2013) and the single protein FusA (Ambroset et al., 2017). The monophyletic clustering of all *M. feriruminatoris* strains is also in agreement with the definition of a homogeneous species. The global phylogeny therefore indicates that *M. feriruminatoris* is more closely related to the *M. mycoides* cluster than to *M. putrefaciens* and *M. yeatsii.*

**Figure 5.**
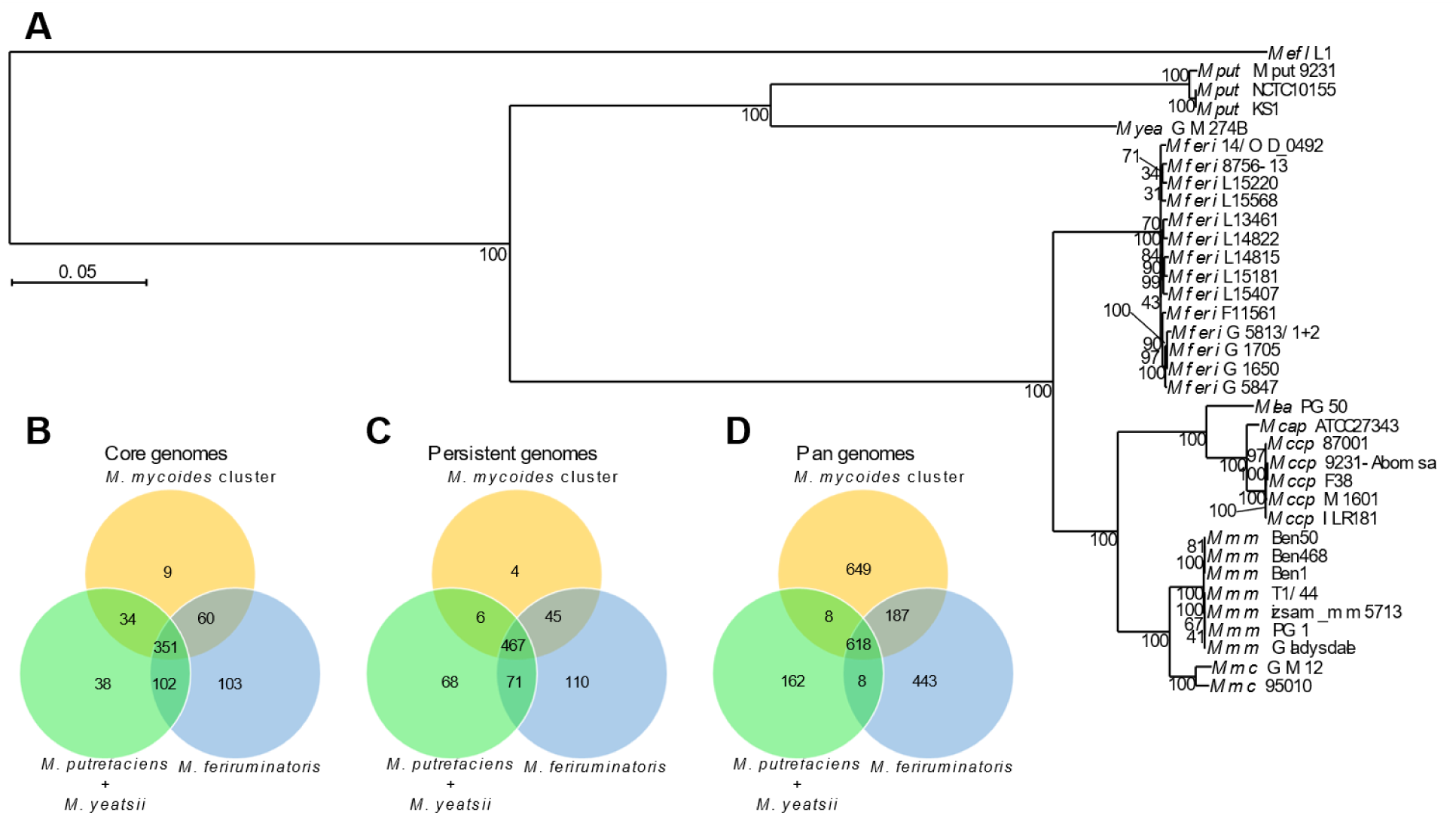
Comparison between *M. feriruminatoris*, the *M. mycoides* cluster, and their closest relatives. **(A)** A phylogenetic tree inferred using the maximum-likelihood method from the concatenated alignments of 294 single-copy core protein sequences, resulting in a total of 92,785 amino acid sites in the alignment matrix. *Me. florum* L1 was used as the outgroup, and 150 bootstrap replicates were run. Bootstrap values are shown as node labels. Venn diagrams of the protein clusters from the **(B)** core genome, **(C)** persistent genome, and **(D)** pan-genomes shared between *M. feriruminatoris,* the *M. mycoides* cluster, and their closest relatives.

### What makes the difference between *M. feriruminatoris* and related species?

To identify specificities of the *M. feriruminatoris* species compared to phylogenetically-related ruminant species, core and pan-genomes were constructed for three sets of genomes including 14 *M. feriruminatoris* strains, 16 *M. mycoides*-cluster strains and 4 *M. yeatsii* and *M. putrefaciens* strains, respectively. The core genomes of each set were similar in size, with 616 clusters for *M. feriruminatoris*, 454 for the *M. mycoides*-cluster strains, and 525 for *M. putrefaciens* and *M. yeatsii*. We determined the overlap between these core genomes (Figure 5B) and found that 351 protein clusters were shared by all three core genomes. The *M. feriruminatoris* core genome intersects with a very similar number of genes to the other two, sharing 453 protein clusters with the *M. putrefaciens* and *M. yeatsii* core and 411 with the *M*. *mycoides*-cluster core. The number of shared protein clusters was extended to 467 when considering the persistent genomes instead of the strict core genomes (Figure 5C). We also compared the pan-genomes of the three sets (Figure 5D). The relatively small pan-genome for the *M. putrefaciens* and *M. yeatsii* set (796 protein clusters) might be explained by both their small genome sizes and the limited number of genomes available. In contrast, the pan-genomes of the two other sets were larger, with 1,256 clusters for *M. feriruminatoris* and 1,462 for the *M. mycoides* cluster. As these last two sets were composed of a similar number of ∼1 Mbp genomes (Table S2), this result suggests that the gene diversity within the *M. feriruminatoris* species was comparable to the gene diversity of the whole *M. mycoides* cluster. A total of 618 protein clusters were present in at least one strain of the three sets, which represents 49.2%, 42.3% and 77.6% of the *M. feriruminatoris* pan-genome, *M. mycoides*-cluster pan-genome, and *M. putrefaciens*–*M. yeatsii* pan-genome, respectively. The *M. putrefaciens*–*M. yeatsii* pan-genome barely intersects with the other two sets, sharing only 8 protein clusters with each, which further illustrates the distance between this group and the other two groups that overlap with 187 protein clusters.

To document the evolution of gene repertoires during the speciation process of *M. mycoides* cluster-related ruminant mycoplasmas, we employed an ancestral genome reconstruction approach using the birth-and-death model implemented in COUNT (Csurös, 2010). Starting from the content of the protein clusters and the phylogenetic relationship of included genomes, ancestral genome contents were simulated at each node of the phylogenetic tree with posterior probabilities for the protein cluster sizes (Figure S4). We thus produced aggregate information for each inner node (number of clusters present with posterior probabilities superior to 0.5) and for each edge leading to the node (cluster gains, cluster losses). The last common ancestor (LCA) of *M. feriruminatoris* was proposed to include 805 protein clusters (node 16), which is close to the average calculated from the 14 *M. feriruminatoris* genomes with 786 protein clusters. Evolution from the *M. feriruminatoris*/*M. mycoides* LCA (node 32) to the *M. feriruminatoris* LCA (node 16) was associated with the gain of 17 protein clusters and the loss of 25 protein clusters (Table S3). Among the 17 gained clusters, 7 (41.2%) are specific to the *M. feriruminatoris* core genome and totally absent from species belonging to the *M. mycoides* cluster. Further investigation indicated that three clusters might be involved in bacterial defense mechanisms, and four were predicted as surface proteins.

The 25 lost protein clusters included eight involved in metabolism (mainly sugar transport and metabolism), seven involved in information storage and processing (i.e. restriction modification systems and MGE), and one involved in cellular processes and signaling. Note that some of the predicted losses during *M. feriruminatoris* speciation were also predicted to have happened during the evolution of other caprine mycoplasmas. This was the case for genes involved in trehalose metabolism and for the MurR-RpiR family transcription regulator that were also lost in the *M. putrefaciens/M. yeatsii* LCA and *Mcap/Mccp* LCA but maintained in the *M. leachii* LCA and the *Mmm/Mmc* LCA. The main nodes leading to species (nodes 16, 21, 30) and subspecies (nodes 20 and 28) clearly show that these evolution steps were marked by gains of genes mainly associated with defense systems and surface proteins and by losses of genes involved in various aspects of cellular life, notably carbohydrate metabolism (Table S3). These differences in gene categories involved in gains and losses during the emergence of *M. feriruminatoris* suggest that the speciation process might be associated with key metabolic changes and with developments of elements (surface proteins) involved in host–pathogen interaction.

One of the main traits of the *M. feriruminatoris* species is its fast-growing capacity in axenic media, with a reported doubling time of ∼30 min at 37°C (Jores et al., 2013). In comparison, members of the *M. mycoides* cluster have generation times ranging from 80–200 min (March et al., 2000; Lartigue et al., 2007; Jores et al., 2013; Jores et al., 2019; Hutchison III et al., 2016). Our comparative analyses did not identify specific metabolic pathways that could explain this noticeable difference. In order to further investigate the genetic basis of the fast-growing *M. feriruminatoris* phenotype, we compared the genomic regions encompassing the predicted replication origins of the chromosomes of *M. feriruminatoris* strains and related species (Figure S5). All *M. feriruminatoris oriC* regions were highly similar, with intergenic sequences located upstream and downstream of the *dnaA* gene containing 7 and one predicted DnaA boxes, respectively. Interestingly, upstream of the *dnaA* gene, one DnaA box perfectly matched the optimal consensus TTATCCACA in all *M. feriruminatoris* strains, whereas only imperfect DnaA boxes were predicted in *M*. *mycoides*-cluster species. Downstream of the *dnaA* gene, one perfect DnaA box was found in *M. feriruminatoris* and in all other strains from the *M. mycoides* cluster. This remarkable feature suggests that the replication initiation process might be accelerated by enhanced binding of DnaA to the *M. feriruminatoris oriC* region. Further experiments based on DnaA box mutagenesis will be necessary to test this hypothesis.

In conclusion, this comparative genomics study confirmed the genomic boundaries of the *M. feriruminatoris* species. *M. feriruminatoris* has a closed pan-genome extrapolated from strains collected in different localizations over a 30-year period. The intraspecies variability is limited and mainly due to mobile elements such as IS, MICEs, and even a plasmid detected in one isolate only. The *M. feriruminatoris* species is very closely related to the *M. mycoides* cluster, as demonstrated by its phylogenetic positioning but also by its gene content and genome organization as well as several typical characteristics (plasmid, lipoprotein repertoire, production of galactan and glucan, etc.). Therefore, we propose to extend the perimeter of the *M. mycoides* cluster to include the *M. feriruminatoris* species. The evolution of *M. feriruminatoris* is associated with both losses and gains of genes, but further studies will be necessary to determine if these could explain the host specificity of *M. feriruminatoris* to wild *Caprinae*. Indeed, no spillover to domesticated ruminants has been detected so far. Finally, yet importantly, further work is needed to assess whether the specific organization and structure of the DnaA boxes around the *oriC* of the *M. feriruminatoris* genomes could explain its growth characteristics. The recent development of highly efficient in-yeast genome engineering methods and genome transplantation protocols for *M. feriruminatoris* (Talenton et al., 2022) now opens up ways to tackle the questions raised by our study.

## Supporting information

Baby 2023 Supplementary Information

Baby 2023 Table S1

Baby 2023 Table S2

Baby 2023 Table S3

## Acknowledgments

The authors thank Pamela Nicholson for her technical help. We thank the team of the Centre de Calcul Scientifique at the Université de Sherbrooke for their valuable technical assistance. Access to computational resources was provided in part by Calcul Québec (http://www.calculquebec.ca) and Compute Canada (http://www.computecanada.ca). We also acknowledge important input from the Vigimyc network, which made it possible to collect most of the French isolates and Alain Blanchard for fruitful discussions on the manuscript. This research was supported by the International Development Research Centre (Grant No. 108625, https://www.idrc.ca).

