## Supplementary material for "Comparative genomics of *Mycoplasma feriruminatoris*, a fast-growing pathogen of wild *Caprinae*": Baby 2023 Supplementary Information

[1] Univ. Bordeaux, INRAE, UMR BFP, F-33882, Villenave d'Ornon, France
[2] Université de Lyon, Anses–Laboratoire de Lyon, VetAgro Sup, UMR Mycoplasmoses animales, 69007 Lyon, France
[3] Department of Biology, University of Fribourg and Swiss Institute of Bioinformatics, CH-1700 Fribourg, Switzerland
[4] Université de Bordeaux, INRAE, BIOGECO, 33610 Cestas, France
[5] Institute of Veterinary Bacteriology, Vetsuisse Faculty, University of Bern, CH-3001 Bern, Switzerland
† These authors contributed equally to this work.

**Supplementary figures.**

A

| COG DOMAIN | eggNOG | CORE | PERSISTENT | ACCESSORY | SINGLETON |
|---|---|---|---|---|---|
| **INFORMATION STORAGE AND PROCESSING** | | | | | |
| Translation, ribosomal structure and biogenesis | J | 109 | 126 | 0 | 27 |
| Transcription | K | 30 | 32 | 6 | 16 |
| Replication, recombination and repair | L | 47 | 54 | 35 | 63 |
| **CELLULAR PROCESSES AND SIGNALING** | | | | | |
| Cell cycle control, cell division, chromosome partitioning | D | 12 | 14 | 5 | 10 |
| Cell wall/membrane/envelope biogenesis | M | 7 | 17 | 2 | 4 |
| Posttranslational modification, protein turnover, chaperones | O | 14 | 14 | 2 | 4 |
| Signal transduction mechanisms | T | 7 | 8 | 0 | 2 |
| Intracellular trafficking, secretion, and vesicular transport | U | 10 | 13 | 1 | 4 |
| Defense mechanisms | V | 13 | 16 | 16 | 22 |
| **METABOLISM** | | | | | |
| Energy production and conversion | C | 27 | 29 | 0 | 4 |
| Amino acid transport and metabolism | E | 24 | 26 | 4 | 8 |
| Nucleotide transport and metabolism | F | 36 | 38 | 0 | 3 |
| Carbohydrate transport and metabolism | G | 41 | 45 | 8 | 0 |
| Coenzyme transport and metabolism | H | 16 | 20 | 6 | 11 |
| Lipid transport and metabolism | I | 17 | 17 | 1 | 3 |
| Inorganic ion transport and metabolism | P | 33 | 34 | 3 | 8 |
| Secondary metabolites biosynthesis, transport and catabolism | Q | 1 | 3 | 0 | 0 |
| **UNKNOWN** | | | | | |
| Function unknown | S | 87 | 94 | 52 | 48 |
| Not assigned | no | 99 | 121 | 92 | 132 |
| TOTAL | | 630 | 721 | 233 | 369 |

B

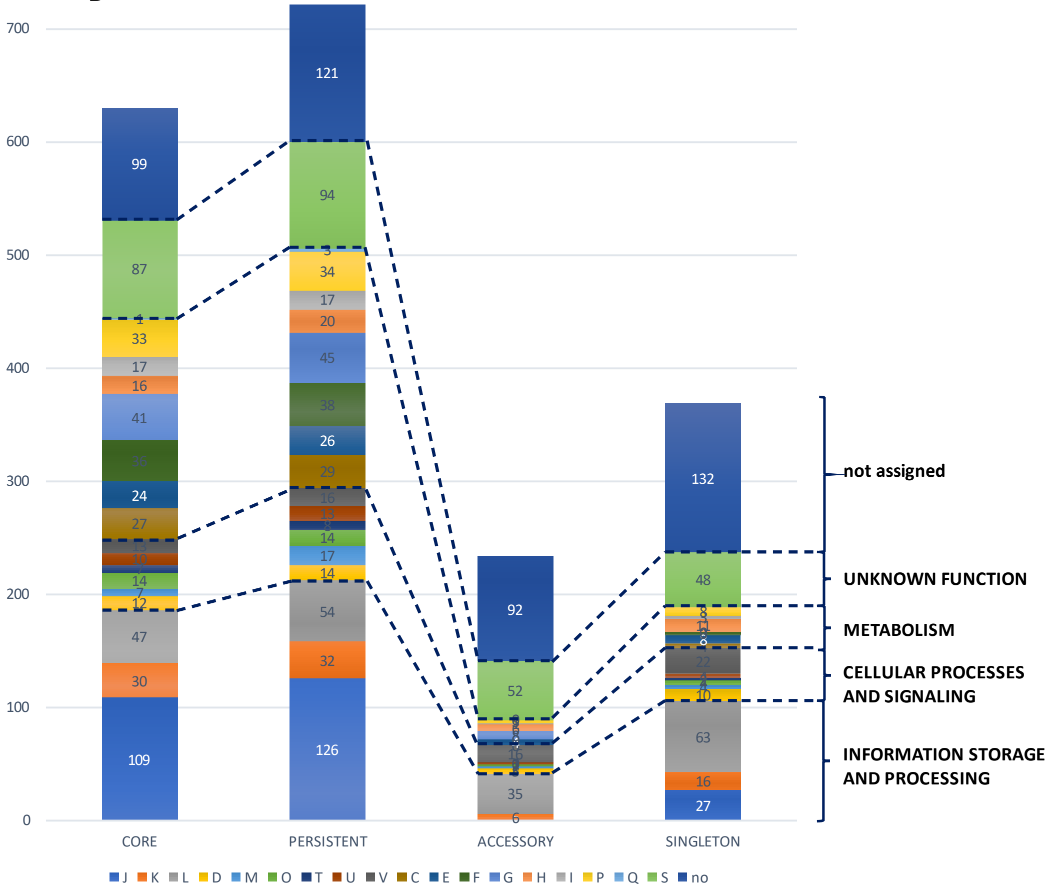


**Figure S1.** Distribution of pan -genome gene families into function EggNOG categories. **(A)** Table showing EggNOG and COG domain distribution and the number of genes associated to each category. **(B)** For *M. feriruminatoris* core, persistent, accessory and singleton genomes, the distribution of gene families into non-supervised orthologous groups (NOGs shown as capital letters) is represented by a diagram. Number of gene families are indicated. Gene families with no EggNOG assignment are not considered.

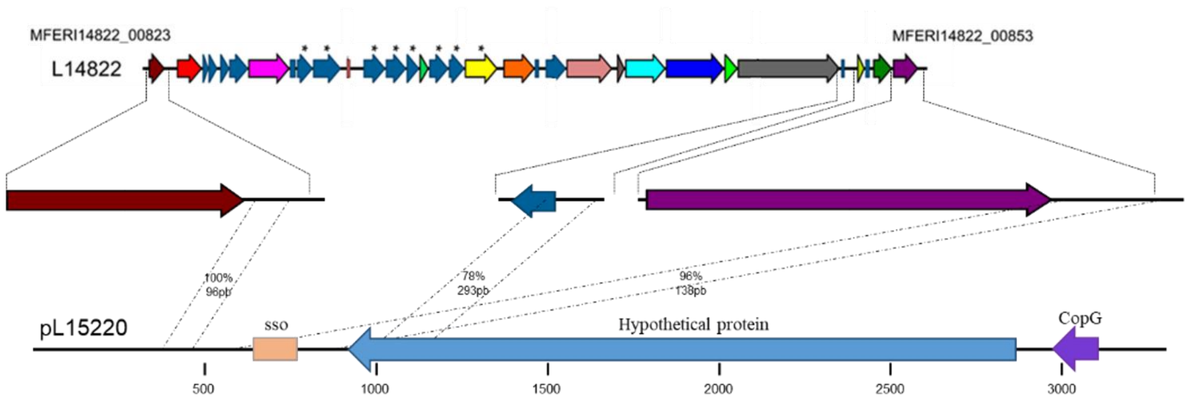


**Figure S2.** Schematic representation of the pL 15220 plasmid and homolog sequences in the L 14822 MICE. The plasmid CDS encodes 2 ORFs ; CopG (purple arrow) and an hypothetical protein (blue arrow). The single strand origin sso is represented by a pink box . Dotted lines highlight sequences with 78 to 100 % nucleotide identity between MICE and the plasmid.

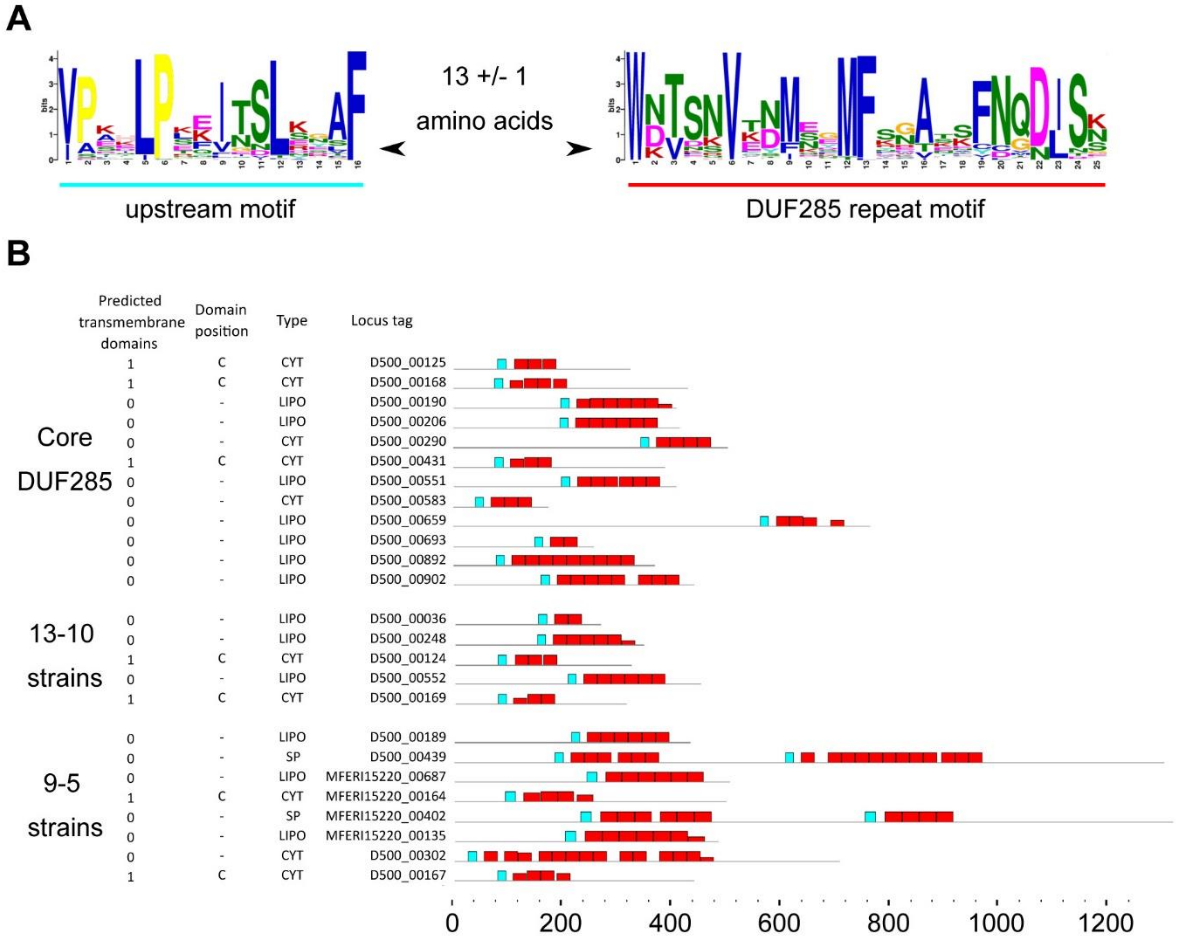


**Figure S3**. DUF 285 family proteins in *M. feriruminatoris*. **(A)** Logos of the DUF 285 repeat, its preceding motif and the average distance in between. The colored rectangle below the logos corresponds to the color used in B to depict the motifs. **(B)** Variability of the DUF 285 family of proteins in the *M. feriruminatoris* strains. A single representative of the most conserved clusters of DUF 285 family of proteins is shown. The proteins are classified based on their level of conservation. The number of predicted transmembrane domains and their relative location on the protein sequence (either close to the C - , N - or both termini) is shown. The predicted localization of the protein is represented by either CYT for cytosolic, LIPO for lipoprotein or SPI for membrane proteins with a SPI type signal peptide. The name of the representative sequence as well as the location of both the DUF 285 repeats and their preceding motif are represented. The color of the bars indicates the motifs and their height represents the level of identity with the logo.

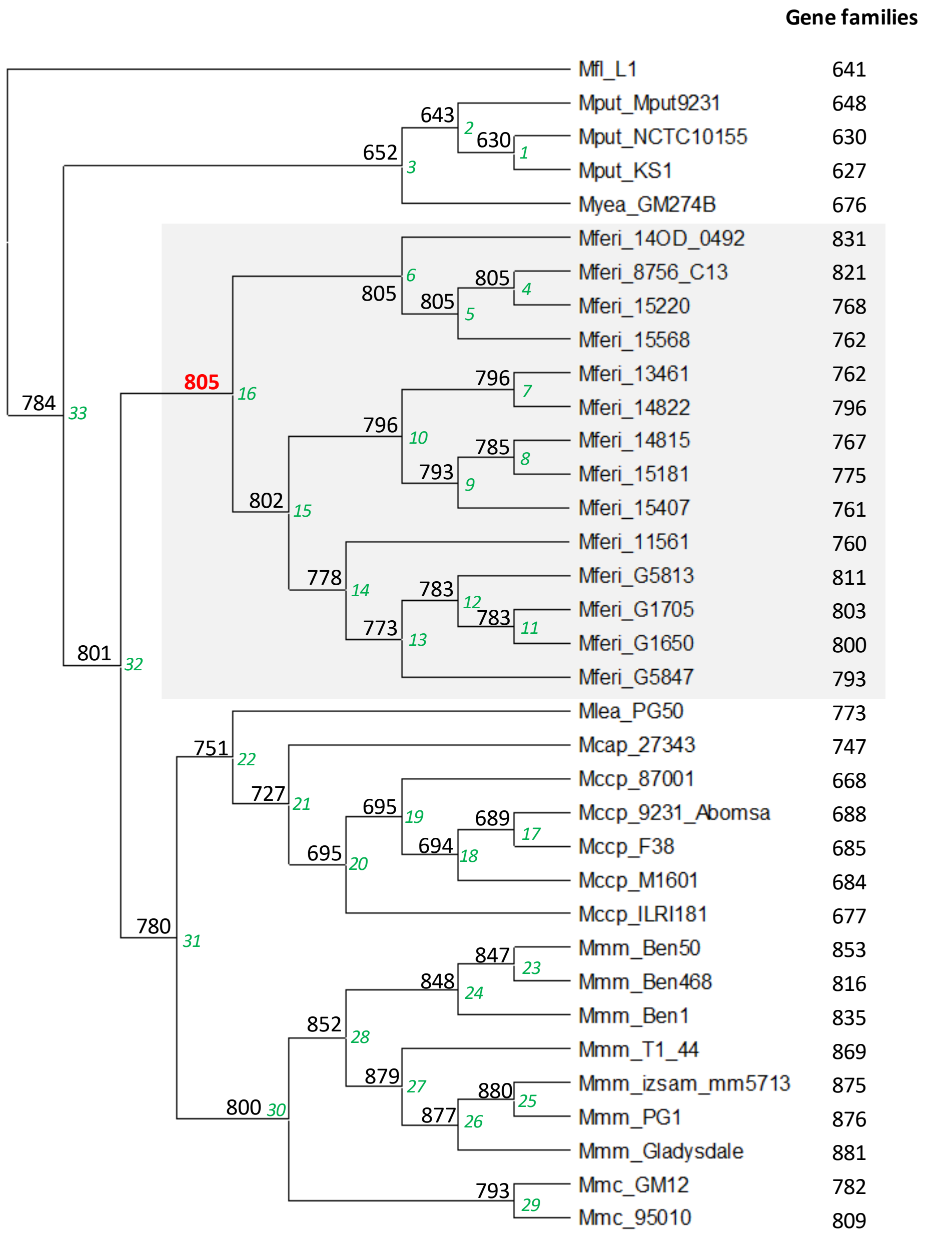


**Figure S4.** Ancestral genome reconstruction. Gene families present at each node of the tree were inferred using the birth - and - death model implemented in COUNT. Nodes are numbered in green, italics. The number of gene families predicted in each ancestral genome is indicated above the corresponding node.

**A-**

```
                        rpmH
MleaPG50-up             CATATCACTTACCTCCTTTACAAAAAAATAA-ATAATTCACACCTAATTATATAATATTT
McapCK-up               CATATCACTTACCTCCTTTACAAAAAAATAA-ATAATTCACACCTAATTATATAATATTT
MccpAbomsa-up           CATATCACTTAGCTCCTTTACAAAAAAATAA-ATAATTCACACCTAATTATATAATATTT
MferiG5813/1+2-up       CATAATACTTACCTCCTTTACAAAAAAATAACATAATTCACACCTAATTATATAATATTT
MferiG1650-up           CATAATACTTACCTCCTTTACAAAAAAATAACATAATTCACACCTAATTATATAATATTT
MferiG1705-up           CATAATACTTACCTCCTTTACAAAAAAATAACATAATTCACACCTAATTATATAATATTT
MferiG5847-up           CATAATACTTACCTCCTTTACAAAAAAATAACATAATTCACACCTAATTATATAATATTT
MferiF11561-up          CATAATACTTACCTCCTTTACAAAAAAATAACATAATTCACACCTAATTATATAATATTT
MferiL15407-up          CATAATACTTACCTCCTTTACAAAAAAATAACATAATTCACACCTAATTATATAATATTT
MferiL15568-up          CATAATACTTACCTCCTTTACAAAAAAATAACATAATTCACACCTAATTATATAATATTT
Mferi8756-C13-up        CATAATACTTACCTCCTTTACAAAAAAATAACATAATTCACACCTAATTATATAATATTT
MferiL13461-up          CATAATACTTACCTCCTTTACAAAAAAATAACATAATTCACACCTAATTATATAATATTT
MferiL14815-up          CATAATACTTACCTCCTTTACAAAAAAATAACATAATTCACACCTAATTATATAATATTT
MferiL14822-up          CATAATACTTACCTCCTTTACAAAAAAATAACATAATTCACACCTAATTATATAATATTT
MferiL15181-up          CATAATACTTACCTCCTTTACAAAAAAATAACATAATTCACACCTAATTATATAATATTT
MferiL15220-up          CATAATACTTACCTCCTTTACAAAAAAATAACATAATTCACACCTAATTATATAATATTT
Mferi14/OD_0492-up      CATAACACTTACCTCCTTTACAAAAAAATAACATAATTCACACCTAATTATATAATATTT
MmcGM12-up              CATACCACT-ACCTCCTTTACAAAAAAATAA-ATAATTCACACCTAATTATATAATATTT
MmmT144-up              CATACCACT-ACCTCCTTTACAAAAAAATAA-ATAATTCACACCTAATTATATAATATTT
                        ****  *** * ******************* ****************************


MleaPG50-up             TGTATTAAAAATACACTTATAAACTTAATTTTTAGCTTCTCTATTAAATGTATATAAAAC
McapCK-up               TGTATTAAAAATACACTTATAAACTTATTTTTTAGCTTATCAATTAAATGTAGATAAAAC
MccpAbomsa-up           TGTATTAAAAATACACTTATAAACTTATTTTTTAGCTTATCAATTAAATGTAGATAAAAC
MferiG5813/1+2-up       TGCTTTAAAAATACACTTATAAACTTAATTTTTAGCTTCTCAATTAAATGTAGATAAAAC
MferiG1650-up           TGCTTTAAAAATACACTTATAAACTTAATTTTTAGCTTCTCAATTAAATGTAGATAAAAC
MferiG1705-up           TGCTTTAAAAATACACTTATAAACTTAATTTTTAGCTTCTCAATTAAATGTAGATAAAAC
MferiG5847-up           TGCTTTAAAAATACACTTATAAACTTAATTTTTAGCTTCTCAATTAAATGTAGATAAAAC
MferiF11561-up          TGCTTTAAAAATACACTTATAAACTTAATTTTTAGCTTCTCAATTAAATGTAGATAAAAC
MferiL15407-up          TGCTTTAAAAATACACTTATAAACTTAATTTTTAGCTTCTCAATTAAATGTAGATAAAAC
MferiL15568-up          TGCTTTAAAAATACACTTATAAACTTAATTTTTAGCTTCTCAATTAAATGTAGATAAAAC
Mferi8756-C13-up        TGCTTTAAAAATACACTTATAAACTTAATTTTTAGCTTCTCAATTAAATGTAGATAAAAC
MferiL13461-up          TGCTTTAAAAATACACTTATAAACTTAATTTTTAGCTTCTCAATTAAATGTAGATAAAAC
MferiL14815-up          TGCTTTAAAAATACACTTATAAACTTAATTTTTAGCTTCTCAATTAAATGTAGATAAAAC
MferiL14822-up          TGCTTTAAAAATACACTTATAAACTTAATTTTTAGCTTCTCAATTAAATGTAGATAAAAC
MferiL15181-up          TGCTTTAAAAATACACTTATAAACTTAATTTTTAGCTTCTCAATTAAATGTAGATAAAAC
MferiL15220-up          TGCTTTAAAAATACACTTATAAACTTAATTTTTAGCTTCTCAATTAAATGTAGATAAAAC
Mferi14/OD_0492-up      TGCTTTAAAAATACACTTATAAACTTAATTTTTAGCTTCTCAATTAAATGTAGATAAAAC
MmcGM12-up              TGCTTTAAAAATACACTTATAAACTTAAATTTTAGCTTCTCAATTAAATGTAGATAAAAC
MmmT144-up              TGCTTTAAAAATACACTTATAGACTTAATTTTTAGCTTCTAAATTAAATGTAGATAAAAC
                        **  ***************** *****  ********* *  ********** *******


MleaPG50-up             TAAAAATTACTCACAAAAATATTAGTTAATTGTTGATAAGTTGATAAGTTATAGATTAGT
McapCK-up               TAAAAATTACTCACAAAAATATTAGTTAATTGTTGATAAGTTGATAAATAATAAATTCAA
MccpAbomsa-up           TAACAATTACTCACAAAAATATTAGTTAATTGTTGATAAGTTGATAAATAATAAATTCAA
MferiG5813/1+2-up       TAGAAATTACTAACAAAAATATTAGCTAATTGTGGATAAGTGAATAAGTTATAAATAACA
MferiG1650-up           TAGAAATTACTAACAAAAATATTAGCTAATTGTGGATAAGTGAATAAGTTATAAATAACA
MferiG1705-up           TAGAAATTACTAACAAAAATATTAGCTAATTGTGGATAAGTGAATAAGTTATAAATAACA
MferiG5847-up           TAGAAATTACTAACAAAAATATTAGCTAATTGTGGATAAGTGAATAAGTTATAAATAACA
MferiF11561-up          TAGAAATTACTAACAAAAATATTAGCTAATTGTGGATAAGTGAATAAGTTATAAATAACA
MferiL15407-up          TAGAAATTACTAACAAAAATATTAGCTAATTGTGGATAAGTGAATAAGTTATAAATAACA
MferiL15568-up          TAGAAATTACTAACAAAAATATTAGCTAATTGTGGATAAGTGAATAAGTTATAAATAACA
Mferi8756-C13-up        TAGAAATTACTAACAAAAATATTAGCTAATTGTGGATAAGTGAATAAGTTATAAATAACA
MferiL13461-up          TAGAAATTACTAACAAAAATATTAGCTAATTGTGGATAAGTGAATAAGTTATAAATAACA
MferiL14815-up          TAGAAATTACTAACAAAAATATTAGCTAATTGTGGATAAGTGAATAAGTTATAAATAACA
MferiL14822-up          TAGAAATTACTAACAAAAATATTAGCTAATTGTGGATAAGTGAATAAGTTATAAATAACA
MferiL15181-up          TAGAAATTACTAACAAAAATATTAGCTAATTGTGGATAAGTGAATAAGTTATAAATAACA
MferiL15220-up          TAGAAATTACTAACAAAAATATTAGCTAATTGTGGATAAGTGAATAAGTTATAAATAACA
Mferi14/OD_0492-up      TAGAAATTACTAACAAAAATATTAGCTAATTGTGGATAAGTGAATAAGTTATAAATAACA
MmcGM12-up              AACAAATTACTAACAAAAATATTAGTTAATTGTTGATAAGTTGATAACTATAAAATAACA
MmmT144-up              CACAAATTACTAACAGAAATATTAGTTAATTGTTGATAAGTTGATAA-CTATAAAATAAA
                         *  ******* *** ********* ******* *******  ****     * *
```

```
MleaPG50-up             GCAAATAATTAAGATTTCCT-TTTTTATATTTTGTTAATATAACTATAATATATGGATTTT
McapCK-up               GGAACTAATTAAGATTTTATATTTTTGCATTTTGTTAATATAACTATAATGTATGGATTTT
MccpAbomsa-up           GGAACTAATTAAGATTTTATATTTTTGCATTTTGTTAATATAACTATAATGTATGGATTTT
MferiG5813/1+2-up       CCTTTTAATTAAGATTTTTAATTTCTAGAAAGTGTTAATATAACTATATTATATGGATTTT
MferiG1650-up           CCTTTTAATTAAGATTTTTAATTTCTAGAAAGTGTTAATATAACTATATTATATGGATTTT
MferiG1705-up           CCTTTTAATTAAGATTTTTAATTTCTAGAAAGTGTTAATATAACTATATTATATGGATTTT
MferiG5847-up           CCTTTTAATTAAGATTTTTAATTTCTAGAAAGTGTTAATATAACTATATTATATGGATTTT
MferiF11561-up          CCTTTTAATTAAGATTTTTAATTTCTAGAAAGTGTTAATATAACTATATTATATGGATTTT
MferiL15407-up          CCTTTTAATTAAGATTTTTAATTTCTAGAAAGTGTTAATATAACTATATTATATGGATTTT
MferiL15568-up          CCTTTTAATTAAGATTTTTAATTTCTAGAAAGTGTTAATATAACTATATTATATGGATTTT
Mferi8756-C13-up        CCTTTTAATTAAGATTTTTAATTTCCAGAAAGTGTTAATATAACTATATTATATGGATTTT
MferiL13461-up          CCTTTTAATTAAGATTTTTAATTTCCAGAAAGTGTTAATATAACTATATTATATGGATTTT
MferiL14815-up          CCTTTTAATTAAGATTTTTAATTTCCAGAAAGTGTTAATATAACTATATTATATGGATTTT
MferiL14822-up          CCTTTTAATTAAGATTTTTAATTTCCAGAAAGTGTTAATATAACTATATTATATGGATTTT
MferiL15181-up          CCTTTTAATTAAGATTTTTAATTTCCAGAAAGTGTTAATATAACTATATTATATGGATTTT
MferiL15220-up          CCTTTTAATTAAGATTTTTAATTTCCAGAAAGTGTTAATATAACTATATTATATGGATTTT
Mferi14/OD_0492-up      CCTTTTAATTAAGATTTTTAATTTCCAGAAAGTGTTAATATAACTATATTATATGGATTTT
MmcGM12-up              CCATTAAAATAAGATTTTAGAATTTGTTAATTTGTTAATATATCTATGTTATAAGGATTTT
MmmT144-up              CCATTAAAATAAGATTTTATAATTTATTAATTTGTTAATATATCTATGTTATAAGGATTTT
                              ** ********     **    *   ********** ****  * ** *******

                                    -35                      -10         SD      dnaA
MleaPG50-up             GTCCACGTTTTCCACATTTTTAACAAGTCTTTAATTTATAATATCTTTGGAGATAAATATG
McapCK-up               CTCCACGTTTTCCACATTTTTAACAAGTCTTTACTTTATAATATTTTTGGAGATAAATATG
MccpAbomsa-up           CTCCACGTTTTCCACATTTTTAACAAGTCTTTACTTTATAATATTTTTGGAGATAAATATG
MferiG5813/1+2-up       TTCCACGTTTTCCACATTTTTAACAAGTGTTTAATTTATAATTTTTTTGGAGATAAATATG
MferiG1650-up           TTCCACGTTTTCCACATTTTTAACAAGTGTTTAATTTATAATTTTTTTGGAGATAAATATG
MferiG1705-up           TTCCACGTTTTCCACATTTTTAACAAGTGTTTAATTTATAATTTTTTTGGAGATAAATATG
MferiG5847-up           TTCCACGTTTTCCACATTTTTAACAAGTGTTTAATTTATAATATTTTTGGAGATAAATATG
MferiF11561-up          TTCCACGTTTTCCACATTTTTAACAAGTGTTTAATTTATAATATTTTTGGAGATAAATATG
MferiL15407-up          TTCCACGTTTTCCACATTTTTAACAAGTGTTTAATTTATAATATTTTTGGAGATAAATATG
MferiL15568-up          TTCCACGTTTTCCACATTTTTAACAAGTGTTTAATTTATAATATTTTTGGAGATAAATATG
Mferi8756-C13-up        TTCCACGTTTTCCACATTTTTAACAAGTGTTTAATTTATAATATTTTTGGAGATAAATATG
MferiL13461-up          TTCCACGTTTTCCACATTTTTAACAAGTGTTTAATTTATAATATTTTTGGAGATAAATATG
MferiL14815-up          TTCCACGTTTTCCACATTTTTAACAAGTGTTTAATTTATAATATTTTTGGAGATAAATATG
MferiL14822-up          TTCCACGTTTTCCACATTTTTAACAAGTGTTTAATTTATAATATTTTTGGAGATAAATATG
MferiL15181-up          TTCCACGTTTTCCACATTTTTAACAAGTGTTTAATTTATAATATTTTTGGAGATAAATATG
MferiL15220-up          TTCCACGTTTTCCACATTTTTAACAAGTGTTTAATTTATAATATTTTTGGAGATAAATATG
Mferi14/OD_0492-up      TTCCACGTTTTCCACATTTTTAACAAGTGTTTAATTTATAATATTTTTGGAGATAAATATG
MmcGM12-up              TTCCACGTTTTCCACATTTTTAACAAGTGTTTAA-CTATAATATTTTTGGAGACAAATATG
MmmT144-up              TTCCACGTTTTCCACATTTTTAATAAGTGTTTAA-CTATAATATTTTTGGAGACAAATATG
                         ********************** **** ****   ****** * ******** *******
```

**B-**

```
                          dnaA
MleaPG50-down             TAATAAAAATAACTATTTTAAATCTATGTTTTTATAAGTTATTCACAAATTAACCTCATA
McapCK-down               TAATAAAAATAACTATTTTAAATCTATGTTTTTATAAGTTGTTCACAAATTAACCTCATA
MccpAbomsa-down           TAATAAAAATAACTATTTTAAATCTATGTTTTTATAAGTTGTTCACAAATTAACCTCATA
Mferi/OD_049214-down      TAATAAAA--AATATTTATTAATTTAGGTTTTGTGAGTTATCCACAAATTAACTTCATA
MferiL15220-down          TAATAAAA--AATCTTTATTATTTTAGGTTTTGTGAGTTATCCACAAATTAACTTCATA
MferiG5847-down           TAATAAAA--AATCTTTATTAATTTAGGTTTTGTAAGTTATCCACAAATTAACTTCATA
MferiG5813/1+2-down       TAATAAAA--AATCTTTATTAATTTAGGTTTTGTAAGTTATCCACAAATTAACTTCATA
MferiG1650-down           TAATAAAA--AATCTTTATTAATTTAGGTTTTGTAAGTTATCCACAAATTAACTTCATA
MferiG1705-down           TAATAAAA--AATCTTTATTAATTTAGGTTTTGTAAGTTATCCACAAATTAACTTCATA
MferiF11561-down          TAATAAAA--AATCTTTATTAATTTAGGTTTTGTAAGTTATCCACAAATTAACTTCATA
MferiL14822-down          TAATAAAA--AATCTTTATTAATTTAGGTTTTGTGAGTTATCCACAAATTAACTTCATA
Mferi8756-C13-down        TAATAAAA--AATCTTTATTAATTTAGGTTTTGTGAGTTATCCACAAATTAACTTCATA
MferiL13461-down          TAATAAAA--AATCTTTATTAATTTAGGTTTTGTGAGTTATCCACAAATTAACTTCATA
MferiL14815-down          TAATAAAA--AATCTTTATTAATTTAGGTTTTGTGAGTTATCCACAAATTAACTTCATA
MferiL15181-down          TAATAAAA--AATCTTTATTAATTTAGGTTTTGTGAGTTATCCACAAATTAACTTCATA
MferiL15407-down          TAATAAAA--AATCTTTATTAATTTAGGTTTTGTGAGTTATCCACAAATTAACTTCATA
MferiL15568-down          TAATAAAA--AATCTTTATTAATTTAGGTTTTGTGAGTTATCCACAAATTAACTTCATA
MmcGM12-down              TAATAAAAATAGCTATT-TAAACCTAGATTATTAACAGTTATCCACAAATTAACCTCATA
MmmT144-down              TAACAAAAATAGCAATT-TAAATCTAACTTATTAACAGTTATCCACAAATTAACCTCATA
                          *** ****  *    ** * *   **  ** **   **** * *********** *****

MleaPG50-down             ATAAGAATAATATTTTAGAAATATAATAAAGA------AATAGAAATACAAAATACATCT
McapCK-down               ATAAGAATAATATTTTAGAAATAAATTATAGA------AATAGAAATACAAAACATTCCT
MccpAbomsa-down           ATAAGAATAATACTTTAGAAATAAATTATAGA------AATAGAAATACAAAACATTCCT
Mferi/OD_049214-down      CTAATAATAA---TTTAGAAATAATAT-TAGA------AATAGTAATATAATACAATC--
MferiL15220-down          ATAATAATAA---TTTAGAAATAATAT-TAGA------AATAGTAATATAATACAATC--
MferiG5847-down           CTAATAATAA---TTTAGAAATAATAT-TAGA------AATAGTAATATAATACAATC--
MferiG5813/1+2-down       CTAATAATAA---TTTAGAAATAATAT-TAGA------AATAGTAATATAATACAATC--
MferiG1650-down           CTAATAATAA---TTTAGAAATAATAT-TAGA------AATAGTAATATAATACAATC--
MferiG1705-down           CTAATAATAA---TTTAGAAATAATAT-TAGA------AATAGTAATATAATACAATC--
MferiF11561-down          CTAATAATAA---TTTAGAAATAATAT-TAGA------AATAGTAATATAATACAATC--
MferiL14822-down          CTAATAATAA---TTTAGAAATAATAT-TAGA------AATAGTAATATAATACAATT--
Mferi8756-C13-down        CTAATAATAA---TTTAGAAATAATAT-TAGA------AATAGTAATATAATACAATC--
MferiL13461-down          CTAATAATAA---TTTAGAAATAATAT-TAGA------AATAGTAATATAATACAATC--
MferiL14815-down          CTAATAATAA---TTTAGAAATAATAT-TAGA------AATAGTAATATAATACAATC--
MferiL15181-down          CTAATAATAA---TTTAGAAATAATAT-TAGA------AATAGTAATATAATACAATC--
MferiL15407-down          CTAATAATAA---TTTAGAAATAATAT-TAGA------AATAGTAATATAATACAATC--
MferiL15568-down          CTAATAATAA---TTTAGAAATAATAT-TAGA------AATAGTAATATAATACAATC--
MmcGM12-down              ATATTAATAA---TTTAGAAATAATAT-TAGA------AATAGTAATATATAACAAACCC
MmmT144-down              TTACTAATAA---TTTAGAAATAGAAA-TAGAAATAGTAATAGTAATATATAACAAACCC
                           **  *****   **********      ***      ***** **** *  * *

                                                                              dnaN
MleaPG50-down             TATTTTAATTTATTCTAAATTAAAATAAAAAA---ACATCTAAAAGGAGTAATTATG
McapCK-down               TAATTTAATTTAAATTAAATTAGAATAAAAAACTTATCT-TAAAAGGAGTAATTATG
MccpAbomsa-down           TAATTTAATTTAAATTAAATTAGAATAAAAAACTTATAT-TAAAAGGAGTAATTATG
Mferi/OD_049214-down      ---TCTTATTTATATAAAATAAACTTAGAGAA---------AAAAGGAGTAGATATG
MferiL15220-down          ---TCTTATTTATATAAAATAAACTTAGAGAA---------AAAAGGAGTAGTTATG
MferiG5847-down           ---TCTTATTTATATAAAATAAACTTAGAGAA---------AAAAGGAGTAGTTATG
MferiG5813/1+2-down       ---TCTTATTTATATAAAATAAACTTAGAGAA---------AAAAGGAGTAGTTATG
MferiG1650-down           ---TCTTATTTATATAAAATAAACTTAGAGAA---------AAAAGGAGTAGTTATG
MferiG1705-down           ---TCTTATTTATATAAAATAAACTTAGAGAA---------AAAAGGAGTAGTTATG
MferiF11561-down          ---TCTTATTTATATAAAATAAACTTAGAGAA---------AAAAGGAGTAGTTATG
MferiL14822-down          ---TCTTATTTATATAAAATAAACTTAGAGAA---------AAAAGGAGTAGTTATG
Mferi8756-C13-down        ---TCTTATTTATATAAAATAAACTTAGAGAA---------AAAAGGAGTAGTTATG
MferiL13461-down          ---TCTTATTTATATAAAATAAACTTAGAGAA---------AAAAGGAGTAGTTATG
MferiL14815-down          ---TCTTATTTATATAAAATAAACTTAGAGAA---------AAAAGGAGTAGTTATG
MferiL15181-down          ---TCTTATTTATATAAAATAAACTTAGAGAA---------AAAAGGAGTAGTTATG
MferiL15407-down          ---TCTTATTTATATAAAATAAACTTAGAGAA---------AAAAGGAGTAGTTATG
MferiL15568-down          ---TCTTATTTATATAAAATAAACTTAGAGAA---------AAAAGGAGTAGTTATG
MmcGM12-down              AATTATTTTCTAAAATAAGGTAA---AAACAATTTGTTTTAAAAAGGAGTAATTATG
MmmT144-down              AATTATTTTCTAAAATAAAGTAA---AAACAATTTGATT--AAAAGGAGTAATTATG
                             * *  * **    **   *    * * **         **********  ****
```

**Figure S5.** Alignment of the non-coding sequences located upstream (**A**) and downstream (**B**) from the *dnaA* gene of all 14 *M. feriruminatoris* strains and related species from the mycoides cluster: *M. leachii* strain PG50 (MleaPG50), *Mcap* strain CK (McapCK), *Mccp* strain Abomsa (MccpAbomsa), *Mmc* strain

GM12 (MmcGM12) and *Mmm* strain T144 (MmmT144). Putative DnaA boxes are colored according to their matching score to the DnaA box consensus (TTATCCACA): red, 9/9; green, 8/9; yellow, 7/9. Putative promoter sequences (–35, –10), Shine–Dalgarno (SD) are positioned according to Seto *et al*., 1997. Start and stop codons of the flanking genes (*rpmH* and *dnaN*) are in blue. An asterisk notes positions with identical nucleotide in the four sequences.
